# DamageFormer: a damage-aware multimodal deep learning framework for DNA lesion identification from nanopore sequencing

**DOI:** 10.64898/2026.05.14.725245

**Authors:** Qiang Yang, Lin Li, Qin Ma, Rui Yin

**Affiliations:** Department of Health Outcomes and Biomedical Informatics, College of Medicine, University of Florida, Gainesville, FL 32610, USA; Department of Biomedical Informatics, College of Medicine, The Ohio State University, Columbus, OH 43210, USA

## Abstract

**Background:** DNA lesions arise from endogenous metabolism and environmental exposure and are the major drivers of mutagenesis, aging, and cancer development. However, mapping DNA damage at nucleotide resolution remains a technically challenging task. Nanopore sequencing enables direct detection of chemical perturbations through alterations in ionic current signals. Despite this potential, existing computational approaches remain limited in their capacity to generalize across diverse lesion types and to effectively integrate nucleotide sequence context with raw signal information for accurate detection and localization.

**Results:** We presented DamageFormer, a multimodal deep learning framework for detection and localization of DNA lesions using native nanopore sequencing data. Central to this framework is LesionBERT, a damage-aware genomic foundation model built upon DNABERT-2 and enhanced with lesion-focused reconstruction objectives to improve representation of chemically modified bases. DamageFormer integrated LesionBERT with a neural signal model through an adaptive gating mechanism, enabling dynamic weighting of sequence context and nanopore signal evidence. The model was trained using a joint objective that combines prediction, localization, and contrastive alignment losses to promote cross-modal coherence and spatial precision. On an oxidative DNA damage benchmark comprising paired sequence and signal data, DamageFormer achieved an AUROC of 0.99997 for lesion detection and a mean absolute localization error of 0.00439, consistently outperforming state-of-the-art baselines. Model interpretation analyses revealed context-dependent modality weighting that adapts to variation in signal quality and sequence ambiguity. The proposed framework further generalizes to chemically distinct guanine lesions not observed during the training process, demonstrating its robustness and transferability to unseen damage types.

**Conclusions:** Damage-aware biological language modeling combined with adaptive multimodal fusion enables accurate and interpretable identification of DNA lesions from nanopore sequencing data. This framework provides a scalable approach for characterizing genome-wide damage landscapes and illustrates how chemical DNA information can be systematically incorporated into genomic language models. The source code and pretrained models of this work are available at: https://github.com/UF-HOBIYin-Lab/DamageFormer.

## Background

Reactive oxygen species generated from endogenous metabolism and environmental stressors continuously damage genomic DNA, often referred to as DNA lesions, producing diverse chemical lesions that threaten genome stability^1,2^. Among oxidative modifications, 8-oxo-7,8-dihydro-2′-deoxyguanosine (8-oxo-dG) is the most prevalent lesion, arising from the preferential oxidation of guanine due to its low redox potential^3,4^. It is known that 8-oxo-dG mispairs with adenine during replication drives G: C → T: A transversion mutations that have been implicated in aging, neurodegeneration, and cancer development^5,6^. Beyond oxidative stress, alkylation of guanine at the O6 position generates mutagenic adducts formed by endogenous metabolites and environmental carcinogens^7,8^. The repair of these lesions involves specialized DNA glycosylases, such as 8-oxoguanine DNA glycosylase 1 (OGG1) for oxidative damage and O6-methylguanine-DNA methyltransferase (MGMT) for alkylation damage^9,10^, whose dysfunction has been associated with elevated cancer risk^11,12^. Accurate detection and genomic localization of DNA lesions are essential for understanding how damage patterns shape mutational landscapes. Conventional analytical approaches, such as liquid chromatography-mass spectrometry, provide sensitive quantification but sacrifice positional information^13^. Recent genomic approaches have attempted to address this limitation through indirect detection strategies, including antibody-based enrichment (OxiDIP-seq)^14^, chemical labeling with biotin pull-down (OG-seq)^15^, and detection of repair intermediates (Click-code-seq, OGG1-AP-seq)^16,17^. These studies have shown that DNA damage is not randomly distributed but instead exhibits sequence-and chromatin-dependent patterns across the genome. However, because these methods rely on indirect enrichment or enzymatic processing, they remain susceptible to false positives and limited positional resolution. In particular, their dependence on short-read sequencing restricts analysis of repetitive regions, enrichment-based strategies can introduce false positives due to suboptimal antibody specificity or enzymatic reactivity, and most methods lack single-nucleotide precision^18^. Given the extremely low frequency of oxidative lesions, estimated at 1-100 per million guanines^19^, even modest error rates can obscure biologically meaningful patterns.

Nanopore sequencing, commercialized by Oxford Nanopore Technologies (ONT), offers a fundamentally different paradigm by measuring electrical current changes as native DNA molecules pass through protein nanopores^20,21^. Because chemical modifications alter the physical properties of nucleotides, they produce characteristic current signatures distinct from canonical bases, enabling native detection of base modifications without chemical conversion or amplification^22,23^. This principle has enabled deep learning-based identification of epigenetic modifications such as 5-methylcytosine^24,25^. Recent work has demonstrated that 8-oxo-dG detection is feasible using nanopore sequencing by training neural networks on synthetic oligonucleotides containing the modification in known sequence contexts^26^. Similarly, diverse O6-alkylguanine adducts produce distinguishable signal perturbations that enable their localization and discrimination^27^. These studies establish that nanopore technology can capture damage-specific signatures, yet extracting this information reliably across the full diversity of genomic contexts remains challenging. Nanopore current measurements reflect k-mer-level signals influenced by neighboring bases^20,28^, leading to strong sequence-context dependencies that can obscure subtle damage-induced perturbations^23,29^. Signal variability further arises from pore heterogeneity^30^, translocation kinetics^31,32^, basecalling uncertainty^33^, and experimental conditions^34^, complicating generalization beyond curated training sets. Most existing approaches rely on lesion-specific models trained on limited synthetic data^26,35^, hindering their scalability to diverse damage chemistries and genomic environments.

The emergence of transformer-based language models^36,37^ has transformed biomedical research by enabling context-aware representation learning across diverse data modalities^38–42^. In genomics, large-scale pretrained models such as the Nucleotide Transformer^38^ and DNABERT-2^39^ have demonstrated the ability to capture regulatory grammar, long-range dependencies, and sequence-level functional signals directly from raw DNA. In protein biology, models including ESM^40^ and ProtBERT^41^ learn structural and functional representations from amino acid sequences in a self-supervised manner, supporting downstream tasks such as structure prediction, variant effect estimation, and protein design. These advances illustrate that transformer architectures can internalize latent biological rules from primary sequence alone and transfer this knowledge to diverse predictive applications with minimal task-specific supervision. More recently, multimodal biological models have begun integrating complementary inputs, such as sequence, structure, and experimental signals^24,43,44^, to enhance robustness and generalizability across heterogeneous datasets. Despite this progress, existing DNA language models are trained exclusively on canonical nucleotide sequences and lack awareness of chemical modifications that alter both sequence context and biological function. DNA lesions, in particular, introduce context-dependent perturbations^45^ that are not encoded in standard sequence representations, limiting the applicability of current foundation models^39,46^ to damage-aware tasks. Moreover, effective integration of nucleotide context with nanopore-derived electrical signals requires principled strategies to align heterogeneous modalities within a shared representation space, particularly when the target events (e.g., oxidative or alkylation lesions) are rare relative to the overwhelming background of undamaged bases within the data distribution^47,48^. These gaps motivate the development of a biologically informed, damage-aware genomic language model capable of jointly modeling sequence context and signal perturbations to enable accurate and scalable DNA lesion identification and downstream functional interpretation. Importantly, DNA lesions typically occur at very low frequencies relative to the total number of genomic bases. In genome-scale applications, even marginal false positive rates can translate into thousands of erroneous calls, substantially increasing validation burden and masking genuine biological events. Consequently, lesion detection imposes exceptionally stringent performance requirements, i.e., gains in high-precision regimes, particularly at the extreme upper bounds of accuracy and specificity, are not merely incremental statistical improvements but essential for reliable and large-scale deployment.

In this paper, we introduce a multimodal deep learning framework, called DamageFormer, that integrates a damage-aware genomic language model with raw nanopore signal analysis for accurate DNA lesion detection. This framework aims at four core objectives: modeling lesion-associated sequence context, integrating sequence and physical signal information, jointly predicting lesion presence and position at nucleotide resolution, and learning aligned cross-modal representations that identify true damage events. Specifically, we first develop LesionBERT, a damage-aware genomic language model built through a two-stage training strategy. In the first stage, we perform continued self-supervised pretraining of DNABERT-2 on sequences explicitly encoding four chemically grouped classes of guanine lesions (oxidative, small-alkyl, medium-alkyl, and large-bulky) with an asymmetric masking strategy. In the second stage, the pretrained model is fine-tuned using supervised labels for binary lesion detection, adapting the sequence encoder to the downstream task while preserving the damage-aware representations learned during pretraining. Next, we integrate LesionBERT with a CNN-BiLSTM^49^ signal encoder that models raw nanopore current traces at each genomic locus. The encoded sequence and signal embeddings are combined through a learned gating mechanism that adaptively weights modality contributions.

This adaptive fusion enables the model to emphasize sequence context when signal quality is degraded and to prioritize direct physical measurements when lesion signatures are strong, rather than assuming fixed cross-modal importance^50,51^. To enhance cross-modal discrimination, we incorporate a hard-negative-aware contrastive objective that aligns paired sequence-signal embeddings at lesion sites, promoting consistent joint representations of damage across modalities. The overall training objective integrates focal loss for lesion detection, smooth L1 loss for positional localization, and contrastive alignment with task-specific weighting to balance multi-objective optimization. On an independent 8-oxo-dG nanopore benchmark^26^, our framework achieves near-perfect performance of identifying DNA damage (99.828% in accuracy) and single-nucleotide localization precision (mean absolute error 0.439%).

## Results

### Overview of the DamageFormer architecture

The workflow of the proposed DamageFormer is illustrated in **Fig. 1**. To enable joint detection and localization of DNA lesions from heterogeneous sequencing inputs, we designed a multimodal framework that integrates nucleotide sequence context with raw nanopore signal measurements. By combining DNA sequences and their corresponding electrical signals, the model can capture both sequence-dependent susceptibility patterns and physical signatures of chemical modification. As shown in **Fig. 1A**, this framework comprises five main components: (1) a dataset construction module, (2) a sequence encoder, (3) a signal encoder, (4) a gated fusion module, and (5) a multi-task prediction module with cross-modal contrastive learning. We first constructed the dataset shown in **Fig. 1B** using a standardized sequence-signal processing pipeline. Raw nanopore signals were normalized and basecalled using Dorado^52^. Reads were barcode-identified and aligned to reference constructs to determine sequence identity and lesion coordinates, followed by signal-sequence alignment to map current traces to nucleotide positions. The sequence-signal pairs were assembled into labeled samples indicating lesion presence. For reads lacking raw signal data, a sequence-only pipeline aligned basecalled reads to the reference genome using minimap2^53^ to infer damaged positions without barcode matching or signal alignment. Next, the sequence encoder LesionBERT, a damage-aware genomic language model, was created to generate contextualized embeddings capturing both local sequence motifs and long-range dependencies associated with lesion formation (**Fig. 1B**). In parallel, the signal encoder models raw nanopore current traces associated with each genomic locus (**Fig. 1C**). Following established architectures for nanopore-based modification detection^54,55^, this module combined convolutional layers with residual connections to extract local signal patterns and recurrent layers to capture temporal dependencies in current fluctuations. The signal embeddings summarized characteristic deviations associated with damaged and undamaged bases. Then, the outputs from the sequence and signal encoders were integrated through a learned gating mechanism that dynamically weights modality contributions for each input sample (**Fig. 1D**). The fused representation was then fed into prediction heads for damage detection (primary task) and lesion localization identification (auxiliary task). This multi-task design promotes the model to learn representations that are both discriminative and spatially informative. The model was trained with three types of loss functions: the focal loss or binary cross entropy (BCE) loss for the damage prediction, the smooth L1 loss for the position localization prediction constrained by the damage data, and the contrastive loss for the multimodal data interaction (**Fig. 1E**).

**Fig. 1:**
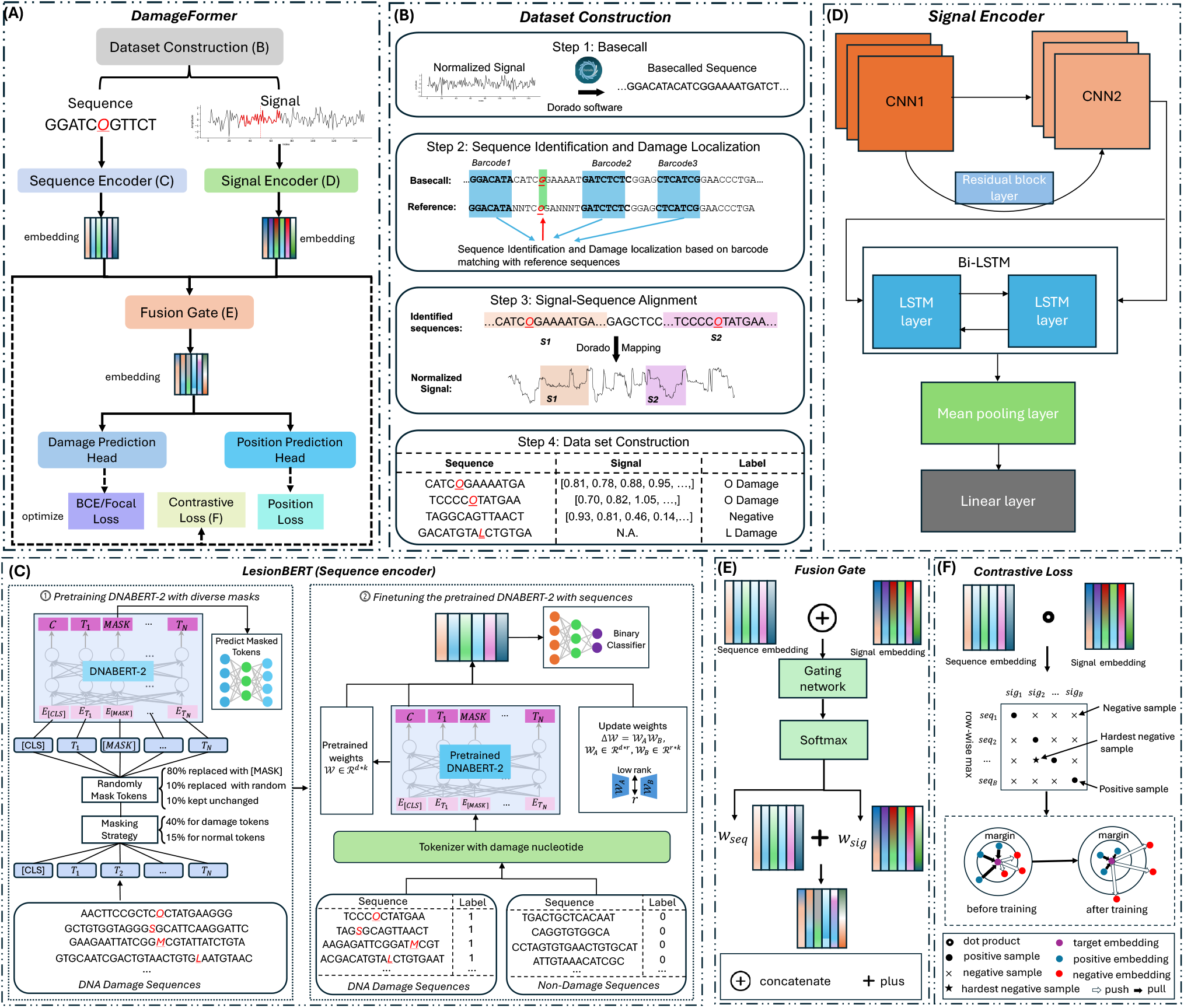
The multimodal deep learning framework for DNA damage detection and localization identification. (A) Overview of the proposed architecture DamageFormer. (B) The workflow of dataset construction for generating labeled sequence-signal pairs from normalized, basecalled, and aligned nanopore reads. (C) The construction of damage-aware foundation genomic language model LesionBERT (sequence encoder) that produces contextualized embeddings, capturing local motifs and long-range genomic dependencies relevant to lesion formation. (D) The main component of signal encoder (CNN-BiLSTM) that extracts hierarchical features from raw nanopore current traces associated with DNA damage. (E) The adaptive fusion gate strategy that dynamically balances sequence and signal representations for each input sample. (F) The hard-negative-aware contrastive learning which aligns matched sequence-signal pairs while encouraging separation between samples with different damage states.

### DNA lesions produce characteristic nanopore signal distortions and structured basecalling errors

Nanopore sequencing records ionic current fluctuations as DNA molecules pass through a protein pore, providing a direct physical readout sensitive to chemical changes in nucleotide structure. To examine how DNA lesions alter this signal landscape, we analyzed raw nanopore traces and basecalling outcomes at ground-truth lesion sites introduced in synthetic oligonucleotides. Inspection of single-molecule reads revealed reproducible local current perturbations surrounding 8-oxo-dG positions (**Fig. 2A**). When aggregated across molecules, these deviations extended beyond the lesion itself and into neighboring nucleotides (**Fig. 2B**), reflecting the multi-base sensing window of the nanopore. Rather than acting as isolated signal spikes, lesion effects propagated across local sequence context, indicating that the observed signal at each position integrates contributions from adjacent bases. This signal propagation was mirrored by systematic decoding instability. Basecalling error rates increased sharply at lesion sites and remained elevated in flanking regions (**Fig. 2C**), revealing a structured error landscape shaped by lesion-induced signal distortion. These patterns suggest that canonical basecalling assumptions break down locally around damaged bases, and that lesion detection must account for distributed contextual effects rather than single-point anomalies. Distinct chemical classes of lesions further produced distinct error profiles (**Fig. 2D**). It is indicated that oxidative, small-alkyl, medium-alkyl, and bulky lesions differed not only in the magnitude of error enrichment at the lesion position but also in the spatial extent of perturbation across neighboring bases. These findings highlight that chemical structure governs both the form and propagation of signal disruption.

**Fig. 2:**
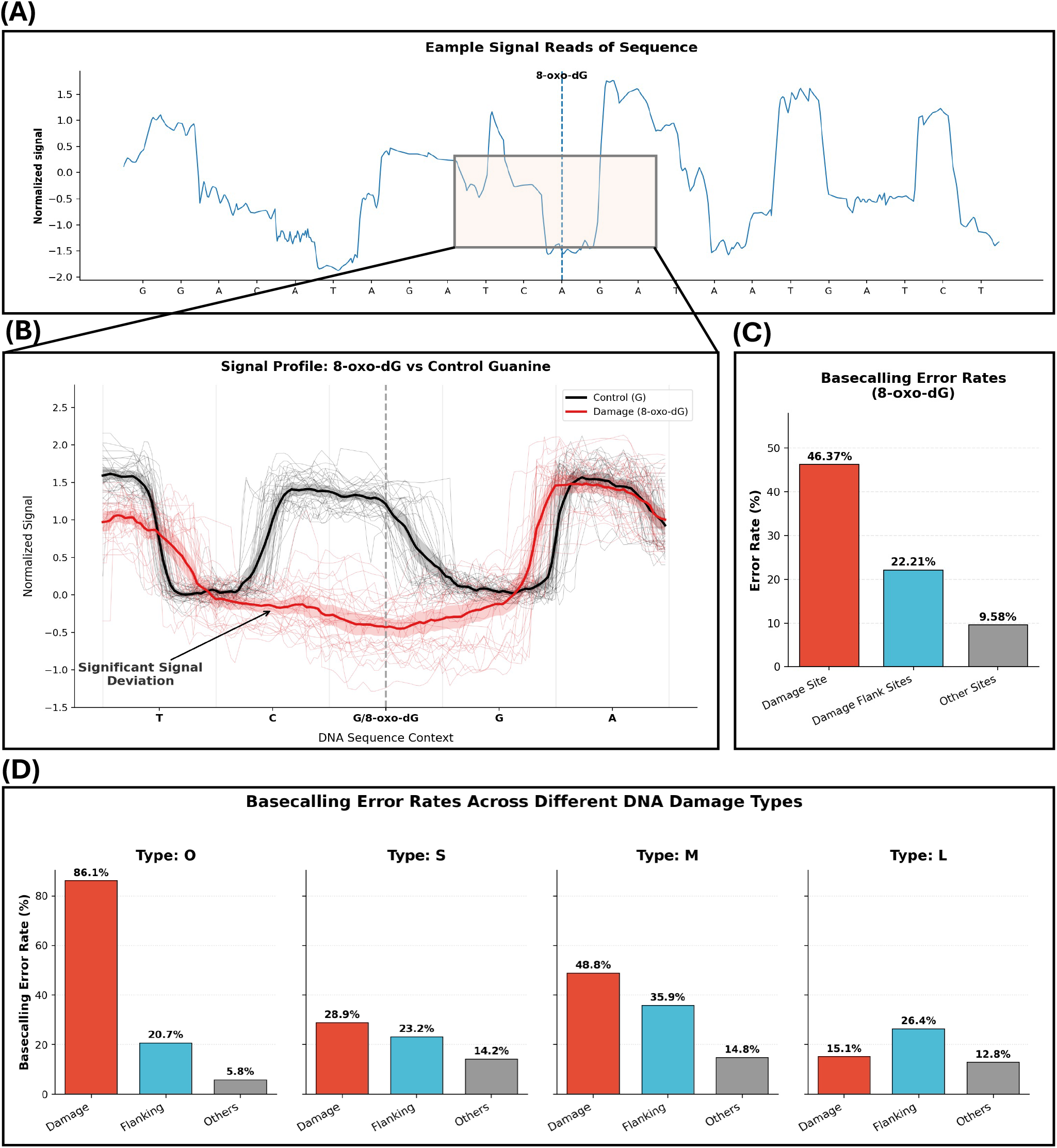
Nanopore signal signatures of DNA damage and their impact on basecalling accuracy. (A) Representative raw nanopore current trace from a DNA molecule containing an 8-oxo-dG lesion. The vertical dashed line marks the lesion position within the nanopore sensing region. (B) Normalized signal profiles for 8-oxo-dG (red) and canonical guanine (black) within identical flanking sequence contexts, illustrating lesion-associated amplitude deviations at and surrounding the modified base. (C) Spatial distribution of basecalling error rates centered on 8-oxo-dG, showing increased errors at the lesion site and neighboring positions relative to distal regions. (D) Basecalling error patterns across four lesion categories (O, S, M, and L), demonstrating damage-type-specific perturbation signatures.

### DamageFormer outperforms existing baseline models

We compared the DamageFormer against a diverse set of baseline methods spanning nanopore-specific deep learning approaches, sequence-based neural models, and traditional machine learning classifiers. Nanopore-based baselines included DeepSignal^24^, NanoCon^43^, Oxo^26^, DeepMod^56^, and Tombo^28^, representing established methods that model nanopore signals with or without sequence context. Sequence-only multilayer perceptron (MLP)-based deep learning baselines included light MLP, deep MLP, and finetuned MLP. Traditional machine learning baselines comprised XGBoost, random forest, linear regression, and support vector machine (SVM). As shown in **Fig. 3A**, the full-scale comparison shows that DamageFormer consistently formed the outer performance boundary, indicating superior results across accuracy, precision, sensitivity, specificity, F1-score, AUROC, and AUPRC. The zoomed panel further confirms that these performance gains remained evident even in the near-ceiling regime. To quantify the contribution of multimodal integration, we compared F1-scores as an example between multimodal models and their signal-only counterparts (**Fig. 3B**). The results suggest that incorporating sequence context yields consistent gains, with an average improvement of +0.022 in F1-score. By contrast, the adaptive gating mechanism in DamageFormer dynamically balances sequence and signal evidence, reducing sensitivity to noisy signal regions and improving discrimination stability. The persistent separation between multimodal and signal-only variants in a near-ceiling regime indicates that these improvements reflect systematic modeling advantages rather than random variation. Similar trends are observed across additional metrics (Additional file 1: **Fig. S1A**).

**Fig. 3:**
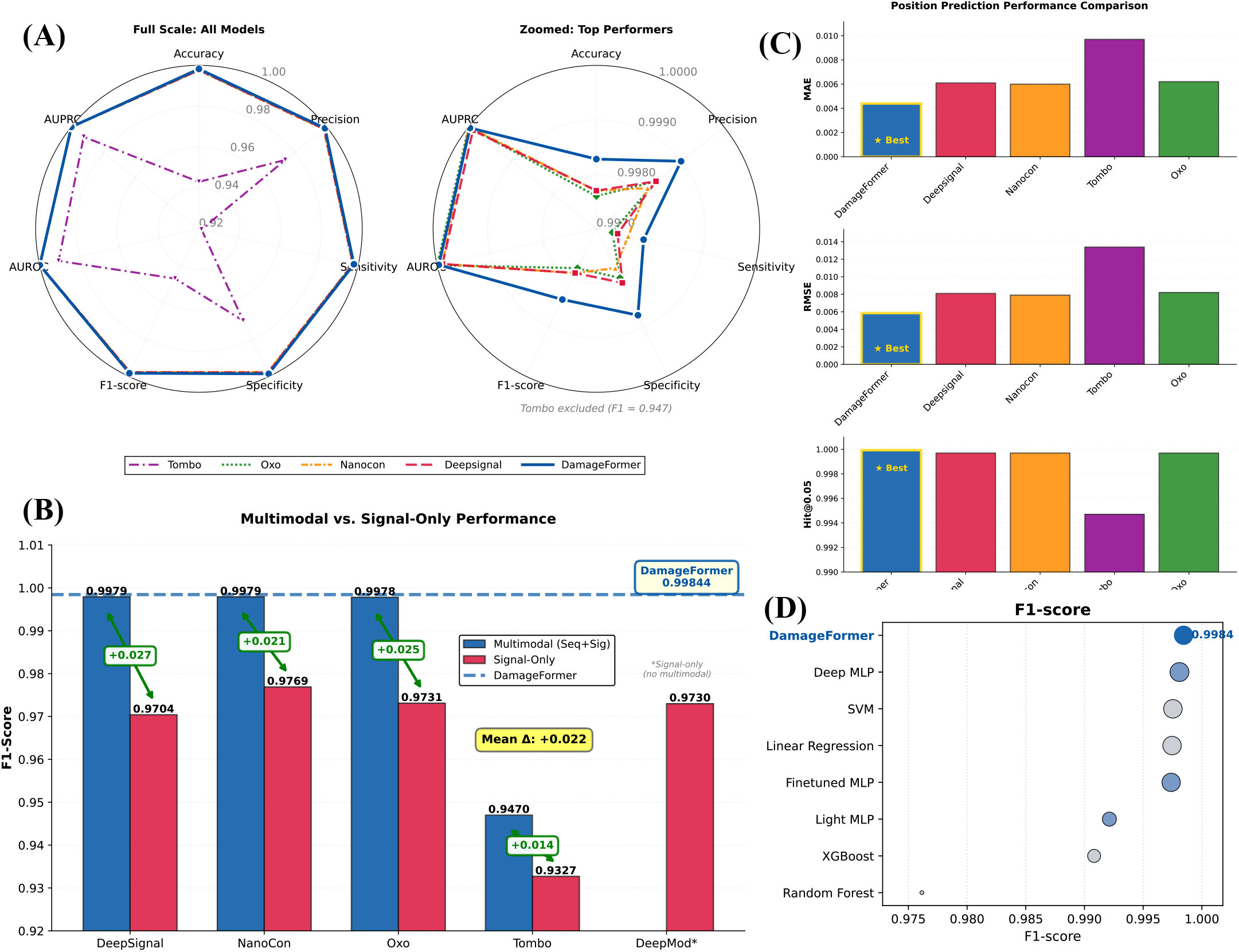
Comprehensive performance comparison of DamageFormer and baseline models. (A) Radar chart summarizing seven detection metrics using both sequence and signal inputs. (B) The F1-scores between multimodal and signal-only variants across architectures. (C) Damage localization prediction performance measured by mean absolute error (MAE), root mean square error (RMSE), and Hit@0.05 accuracy. (D) The F1-score ranking among DamageFormer, nanopore-specific deep learning baselines, sequence-based neural models, and traditional machine learning classifiers.

We further evaluated the performance on the lesion localization task (**Fig. 3C** and Additional file 1: **Fig. S1B**). DamageFormer achieved the lowest mean absolute error (MAE, 0.00439) and root mean square error (RMSE, 0.00585) while simultaneously attaining the highest Hit@0.05 accuracy (0.99993). Performance gains across both error-based and tolerance-based metrics indicate that joint modeling of signal and sequence enhances not only detection confidence but also the accuracy and robustness of lesion localization. The comparison with other baselines further highlighted the advantage of DamageFormer owing to its domain-specific multimodal modeling (**Fig. 3D**). This hierarchy suggests that the improvements stem from principled multimodal representation learning rather than increased model capacity alone. Consistent patterns were also observed across sensitivity, specificity, precision, and accuracy, which further reinforce this interpretation (Additional file 1: **Fig. S1C**).

### LesionBERT promotes DNA lesion prediction and location prediction

To assess the contribution of the LesionBERT in our framework, we conducted systematic comparisons under the same multimodal learning framework across multiple model architectures, focusing on two aspects to examine whether (1) the proposed pretraining and finetuning strategies improve DNA lesion detection and localization prediction, and (2) LesionBERT achieves better performance compared to the original DNABERT-2 and DNABERT models in both tasks.

For the first question, we compared LesionBERT under three regimes: pretraining only, finetuning only, and the full pipeline for DNA lesion and location prediction. As shown in **Fig. 4A**, pretraining alone can provide strong performance, indicating that large-scale sequence modeling could learn transferable structural representations of DNA damage patterns. Finetuning yielded substantial additional improvement, suggesting that task-specific supervision refines these representations toward decision boundaries critical for accurate damage detection. The complete training pipeline further enhanced stability and achieved the most balanced trade-off between sensitivity and specificity, underscoring the complementary roles of both stages. A similar pattern was observed in damage position prediction in **Fig. 4B**. Error-based metrics (MAE and RMSE) decreased sharply after finetuning and reach their minimum in the full model, demonstrating improved localization precision. Meanwhile, tolerance-based localization accuracy (Hit@0.05) increased rapidly after fine-tuning and remained at its peak within the complete pipeline. This pattern indicates that finetuning plays a pivotal role in spatial refinement, while pretraining establishes a robust representational foundation that enables consistent localization.

**Fig. 4:**
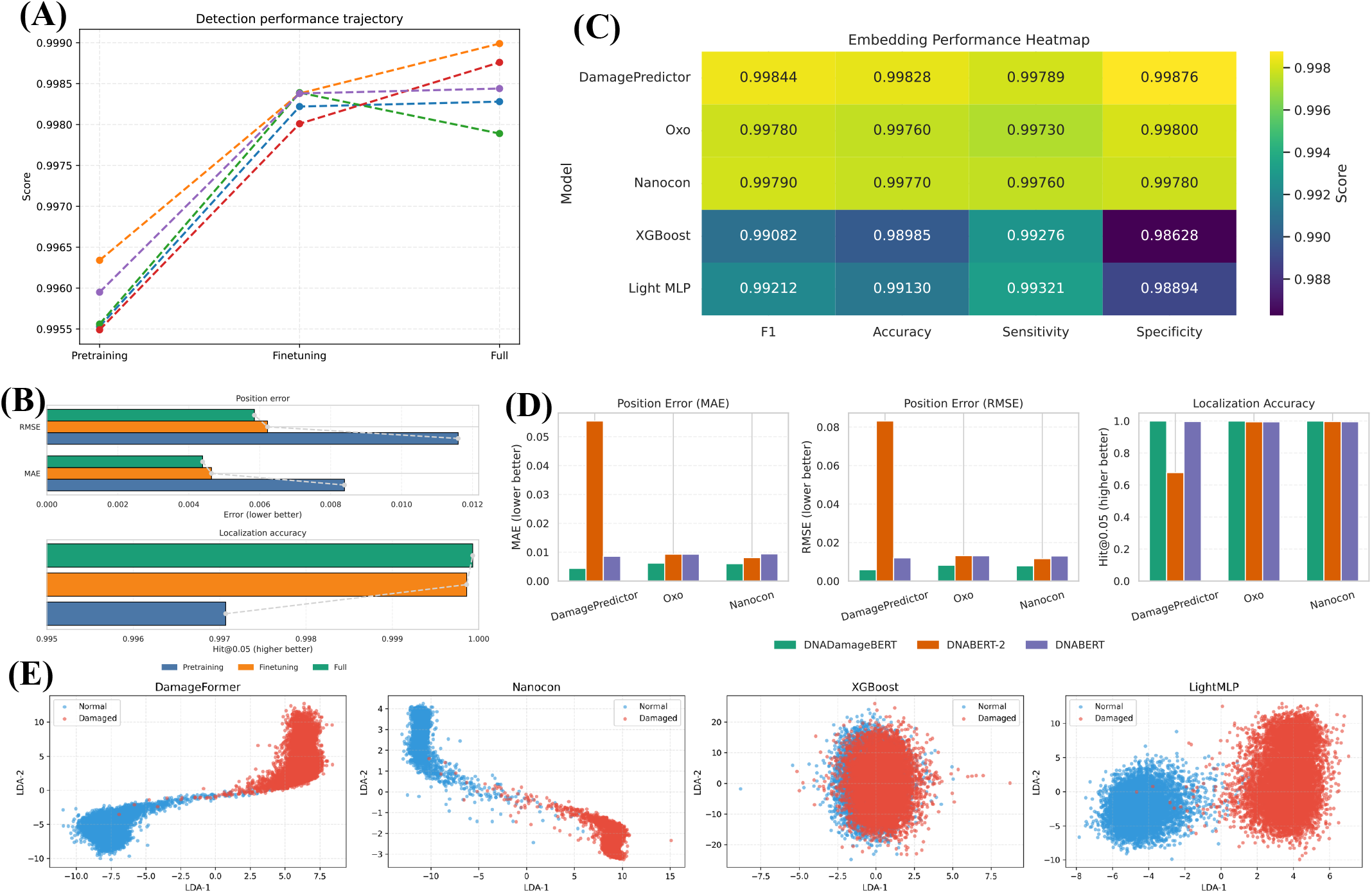
The contribution of LesionBERT to DNA damage detection and localization identification. (A) The DNA damage detection performance across pretraining-only, finetuning-only, and the full training pipeline, evaluated using accuracy, precision, sensitivity, specificity, and F1-score. (B) The performance of damage localization identification measured by mean absolute error (MAE), root mean square error (RMSE) (lower values indicate better performance), and Hit@0.05 localization accuracy (higher values indicate better performance). (C) Heatmap summarizing detection performance (F1-score, accuracy, sensitivity, and specificity) of multiple multimodal architectures under different embedding strategies. (D) Impact of embedding strategies on damage position localization, evaluated by MAE, RMSE, and Hit@0.05 accuracy. (E) Two-dimensional visualization of LesionBERT embeddings for randomly sampled test examples (n = 20,000).

For the second question, we compared LesionBERT with DNABERT-2^39^ and DNABERT^46^ under the same multimodal learning framework across different model architectures, including DamageFormer, Oxo, Nanocon, XGBoost, and Light MLP. As summarized in **Fig. 4C**, LesionBERT consistently achieved the highest or near-highest performance across F1-score, accuracy, sensitivity, and specificity. Performance gains were most pronounced in simpler architectures such as XGBoost and Light MLP, indicating that LesionBERT provided damage-relevant sequence representations not captured by generic nucleotide embeddings. The improvements were balanced across metrics, suggesting enhanced decision boundary stability rather than metric-specific optimization. Moreover, the consistency of these gains across diverse model families supports the conclusion that the performance advantage originates from the embedding space itself. We further evaluated the impact of embedding choice on lesion localization (**Fig. 4D**). Across all architectures, LesionBERT yielded the lowest MAE and RMSE alongside the highest Hit@0.05 accuracy. The improvement is particularly evident in DamageFormer, where DNABERT-2 embeddings resulted in higher positional error, whereas LesionBERT maintained tighter error distributions and consistently high localization accuracy.

To examine how different downstream architectures organize a shared representation space, we visualized embeddings generated by the LesionBERT encoder using supervised dimensionality reduction. Specifically, we combined linear discriminant analysis (LDA)^57^ and principal component analysis (PCA)^58^. Embeddings were projected into a two-dimensional space defined by the LDA axis and an orthogonal PCA component, enabling direct inspection of class separation beyond aggregate performance metrics. As illustrated in **Fig. 4E**, DamageFormer embeddings formed two compact, well-separated clusters, consistent with high linear probe accuracy (0.99750). NanoCon showed slightly weaker but still clear separation (0.99685), whereas XGBoost exhibited substantial class mixing, matching its lower probe accuracy (0.647). The light MLP lies in the middle, producing visible but less compact clusters (0.994). Because these encoders remain fixed across models, these differences reflect how downstream architectures reshape a shared embedding space. The close alignment between geometric cluster separation and probe accuracy indicates that high-performing models reveal class structure directly within the embedding geometry, rather than depending primarily on complex decision boundaries.

### DamageFormer benefits from multimodal data integration and gated fusion

To evaluate the effectiveness of multimodal integration, we compared multimodal, sequence-only, and signal-only configurations under the same training protocol for damage detection. As shown in **Fig. 5A**, the multimodal DamageFormer consistently achieved the highest performance across all metrics, including accuracy, precision, sensitivity, specificity, and F1-score. Sequence-only models retained strong discriminative ability but showed slight reductions in calibration-sensitive metrics, whereas signal-only models exhibited larger declines in sensitivity and balanced accuracy. These results indicated that sequence context and electrical signal features provide complementary information: sequence embeddings captured contextual and structural patterns associated with lesion formation, while signal embeddings encoded local electrophysiological deviations directly induced by damaged bases. The same pattern was observed in positional localization (**Fig. 5B**). The multimodal configuration demonstrated the lowest MAE and RMSE but with the highest Hit@0.05 accuracy, reflecting improvements in both continuous error reduction and tolerance-based correctness. In contrast, sequence-only models provided coarse contextual anchoring, whereas signal-only models captured local fluctuations but lacked global contextual constraints.

**Fig. 5:**
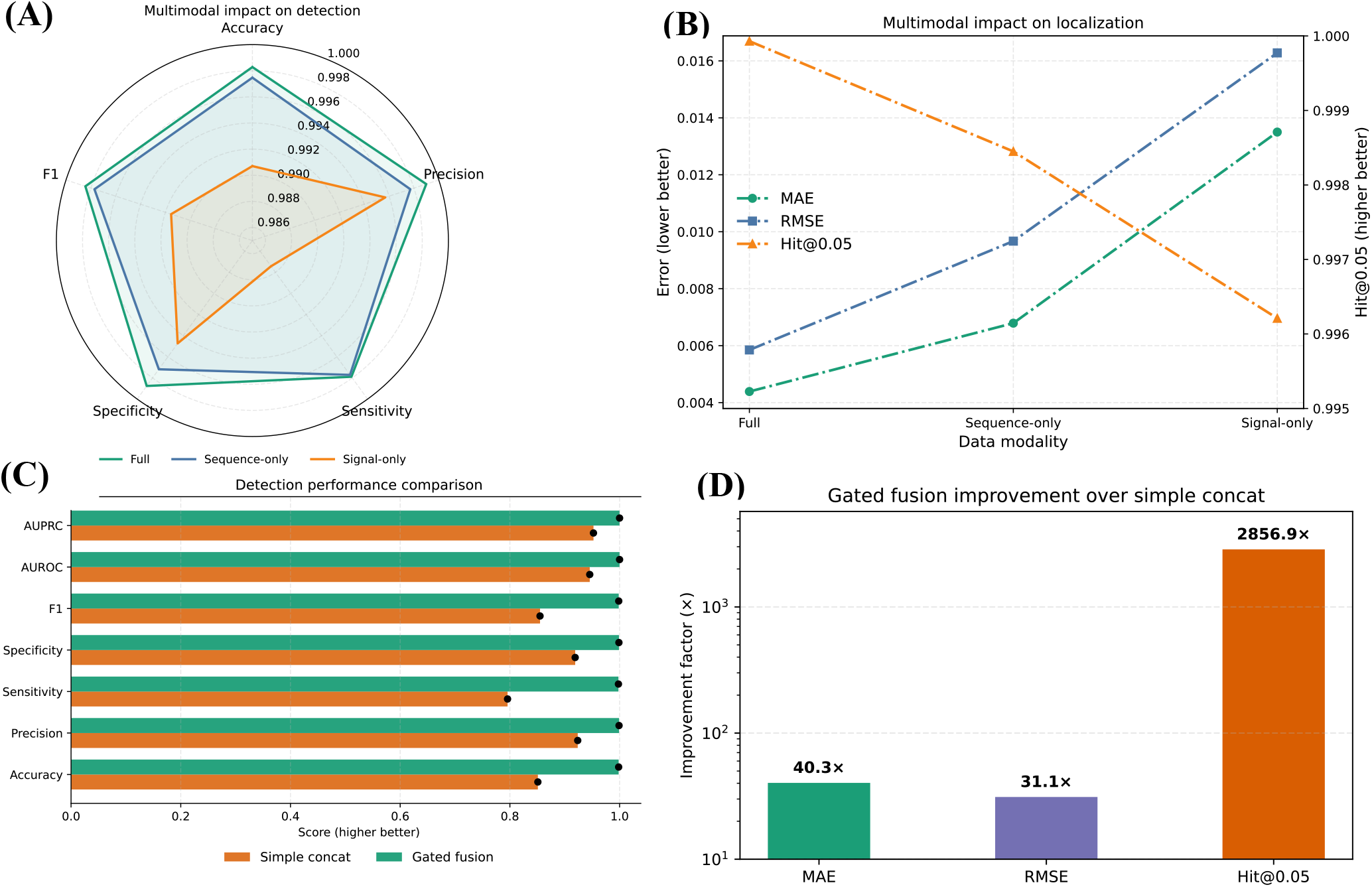
Ablation analysis of multimodal integration and fusion strategies in LesionBERT. (A) Effect of multimodal integration on damage detection performance, comparing full multimodal input (sequence + signal) with sequence-only and signal-only configurations. (B) Effect of multimodal integration on lesion localization accuracy. (C) Detection performance comparison between gated fusion and simple concatenation strategies, evaluated using accuracy, precision, sensitivity, specificity, F1-score, AUROC, and AUPRC. (D) The improvement of gated fusion over simple concatenation for lesion localization. Bars represent fold-change differences (log scale) in MAE, RMSE, and Hit@0.05 accuracy.

To further isolate the role of fusion design, we compared the proposed gated fusion mechanism with a simple concatenation strategy under the identical training conditions. While concatenation treated both modalities equally, gated fusion dynamically reweighted sequence and signal contributions. The results in **Fig. 5C** revealed that gated fusion maintained near-optimal prediction performance, whereas the model with simple concatenation showed systematic degradation, particularly in sensitivity and F1-score. The difference is more pronounced for the localization task (**Fig. 5D**), where gated fusion substantially reduced positional error and markedly improved Hit@0.05 accuracy. These results indicate that adaptive gating is essential for coordinating cross-modal information, acting as a learned arbitration mechanism that can enhance informative signals while suppressing noise.

### Parameter sensitivity and robustness analysis of loss weighting

To evaluate the effect of loss weighting on multimodal training, we systematically varied the coefficients governing positional supervision (*λ*_*pos*_) and contrastive alignment (*λ*_*con*_) while keeping the prediction coefficient, i.e., *λ*_*cls*_, fixed (**Fig. 6**). The positional weight controls the influence of localization error and encourages precise boundary prediction, whereas the contrastive weight regulates cross-modal alignment between sequence and signal embeddings. **Fig. 6A** showed the performance landscape across different coefficient combinations. We observed a broad plateau of near-optimal performance towards damage detection by F1-score and sensitivity, indicating robustness to moderate changes in both weights. In contrast, for damage localization identification, a sharper optimum was shown across MAE and Hit@0.05, highlighting a narrower region where positional accuracy is maximized. The selected configuration (*λ*_*pos*_ = 0.4, *λ*_*con*_ = 0.04) lied within this joint optimum, achieving strong detection performance while minimizing positional error. Increasing *λ*_*con*_ beyond this range produced diminishing returns and may destabilize localization, whereas insufficient positional weighting increases MAE despite preserved prediction performance. These results suggest that balanced auxiliary supervision is essential for stable multimodal learning. To further isolate the influence of contrastive alignment, we fixed *λ*_*pos*_ = 0.4 and varied *λ*_*con*_ (**Fig. 6B**), where we found the results for damage detection remained stable across metrics. Larger contrastive weights could lead to mild sensitivity degradation, indicating that excessive alignment can interfere with prediction calibration, while very small values weaken cross-modal coupling without improving robustness. **Fig. 6C** illustrates the trade-off between detection quality and localization precision across all parameter settings. The result showed that high F1-score and low MAE coexist only within a narrow region, with the selected configuration positioned near the optimal boundary. Parameter settings with larger *λ*_*con*_ tend to deviate from this region, reinforcing the observation that overemphasizing contrastive learning reduces overall efficiency. Finally, robustness analysis across the full parameter grid (**Fig. 6D**) showed that the chosen configuration consistently resided near the favorable edge of the performance distribution, high for detection metrics and low for localization error, rather than representing an unstable outlier or isolated optimum.

**Fig. 6:**
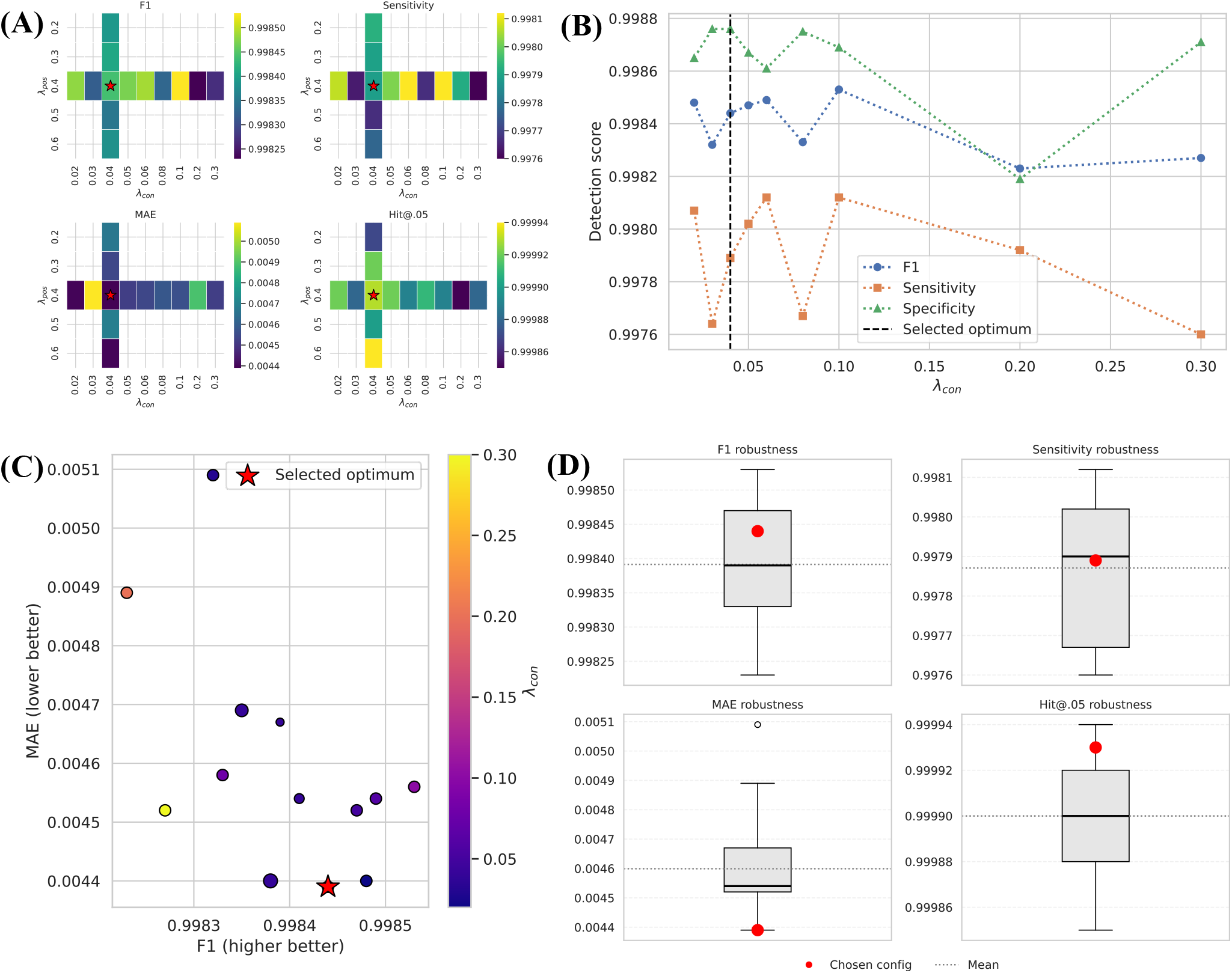
Robustness analysis of loss coefficient weighting in DamageFormer. (A) Performance landscape across combinations of positive-class (damaged) weighting (*λ*_*pos*_) and contrastive loss weighting (*λ*_*con*_). Heatmaps display representative metrics (F1-score, sensitivity, MAE, and Hit@0.05). The red star indicates the selected operating point used in the final model. (B) Parameter sweep showing detection performance as a function of *λ*_*con*_ while fixing *λ*_*pos*_ = 0.4. The dashed vertical line marks the chosen coefficient. (C) Pareto relationship between detection performance (F1-score) and localization precision (MAE). Point color encodes *λ*_*con*_, and marker size represents *λ*_*pos*_. (D) Robustness distributions across all tested coefficient settings. Boxplots summarize variability in F1-score, MAE, sensitivity, and Hit@0.05, with the selected configuration highlighted in red.

### Interpretability of the model’s predictions

To characterize how DamageFormer integrates multimodal information and localizes evidence for DNA damage, we performed complementary interpretability analyses (**Fig. 7**). At the global level, the fusion gate distribution (**Fig. 7A**) showed that the predictions are derived from a wide spectrum of sequence-signal weightings rather than depending consistently on a single modality. To evaluate whether the model’s adaptive fusion weights correspond to meaningful reliance on each modality, we performed cross-modal swap experiments in which either sequence or signal inputs were replaced with those from other samples. Most samples exhibit minimal prediction shifts under both perturbations (**Fig. 7B**). However, sequence swaps produced a broader distribution with heavier tails than signal swaps, indicating that sequence features serve as decisive evidence in a subset of cases, while signal information more often provides complementary support. The strong concentration of shifts near zero further suggests that predictions remained robust under modality perturbation. Consistently, prediction confidence spanned a wide range of fusion weights (**Fig. 7C**). To link these global patterns to individual predictions, we examined representative examples. For a correctly classified damaged sample (sample ID 5604; **Fig. 7D**), both in-silico mutagenesis and gradient-based attribution methods identified a shared localized region with strong influence on model output. The concordance between attribution methods indicates that the model concentrates decision-relevant evidence within a biologically meaningful window rather than dispersing importance across the sequence. In contrast, in a correctly classified undamaged sample (sample ID 55527; **Fig. 7E**), the results showed uniformly weak attribution signals, suggesting that the model predicted the absence of damage based on the lack of strong evidence rather than spurious features.

**Fig. 7:**
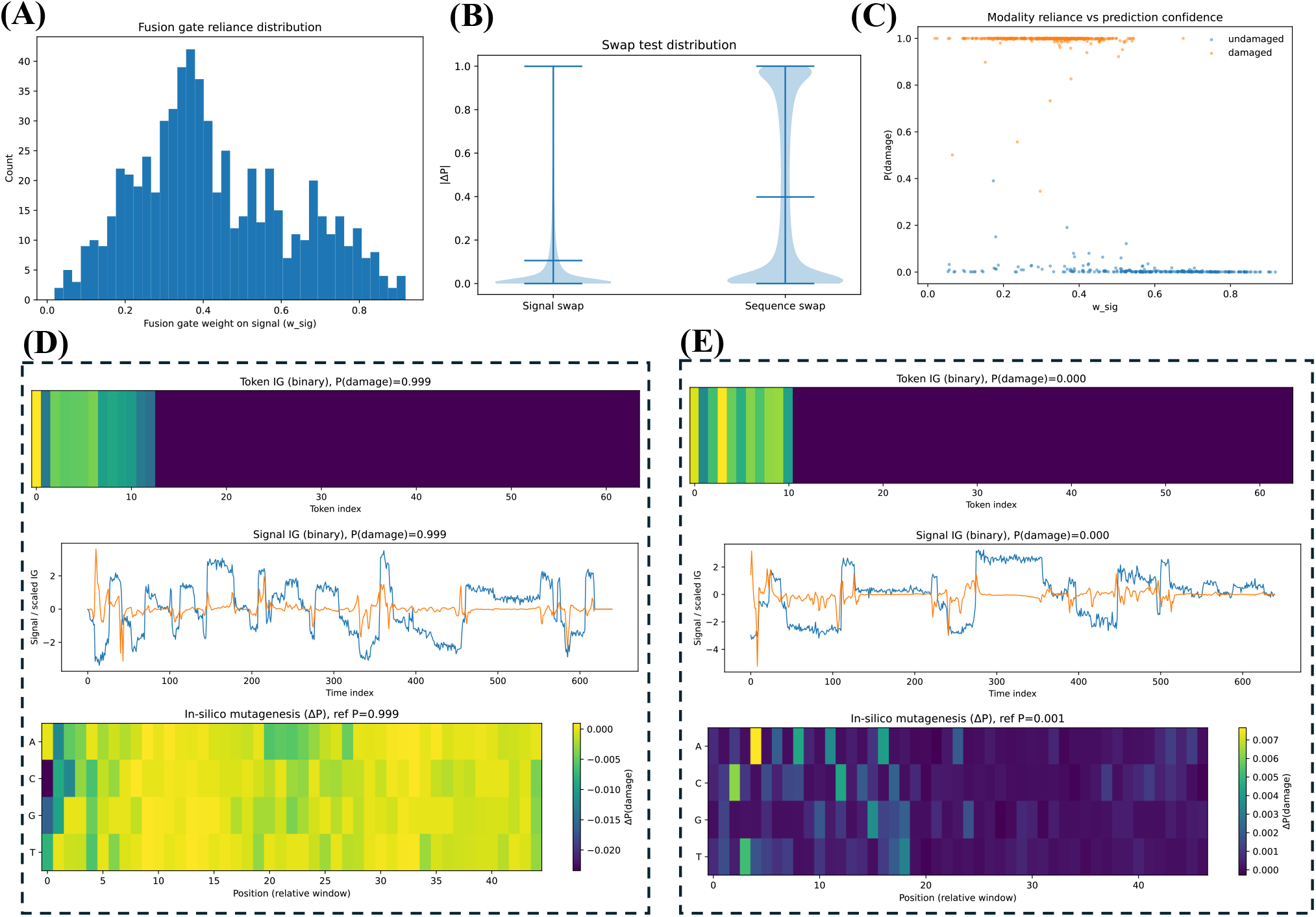
Interpretability analysis of DamageFormer predictions. (A) Distribution of fusion gate weights across the test set, illustrating the relative contributions of sequence and signal modalities. (B) Distribution of absolute prediction shifts (|*ΔP*|) following cross-modal swapping of either signal or sequence inputs. (C) Relationship between modality reliance (fusion weight) and prediction confidence for test samples. (D) Representative damaged sample with attribution analyses, including in-silico mutagenesis, token-level integrated gradients, and signal-based integrated gradients. (E) Representative undamaged sample shown with the same attribution analyses.

### Cross-batch generalization and scalability analysis

To evaluate the scalability and cross-dataset generalization of DamageFormer, we performed a multi-batch analysis using four independently generated batches from a open 8-oxo-dG nanopore dataset^26^, as summarized in **Table 1**. These batches, originally defined in the source study, differ in size and experimental conditions, comprising 8.31%, 19.08%, 35.53%, and 37.08% of the total dataset for Batch 1-4, respectively. Detailed statistics are provided in **Table 2**. We evaluated model performance under two regimes. First, we trained and tested the model independently within each batch (“from-scratch” setting) to assess stability across datasets of varying scale. Second, we trained the model exclusively on Batch 1 and directly applied it to the remaining batches without retraining (“direct-transfer” setting). This design isolates the model’s ability to generalize lesion-aware representations learned from limited data to larger and distributionally distinct datasets. Experimental results show that our DamageFormer maintained strong performance across all batches under both regimes. When trained from scratch, predictive accuracy remained consistently high (≥0.985) despite the large variation in dataset scale, with AUROC and AUPRC values near 1.0 for all batches. Direct transfer from Batch 1 resulted in an expected decline in performance, however, the overall accuracy remained above 0.91 even when the model was evaluated on batches more than four times larger than its training set. A comparable pattern was observed for position localization identification: although error increased modestly under direct transfer, it remained low overall, and Hit@0.05 consistently exceeded 0.997. Notably, the model trained just on 8.31% of the data generalizes effectively to batches comprising up to 37.08% of the total dataset, indicating that the learned representations can capture stable, transferable structure rather than batch-specific artifacts. These results confirm that DamageFormer scales robustly and preserves predictive quality across independently sampled datasets of different sizes.

**Table 1:** Cross-batch evaluation of DamageFormer under two training regimes. (A) DNA damage prediction. (B) Damage position localization. For each batch, results are reported under two settings: (1) training and testing from scratch within the same batch, and (2) direct-transfer evaluation, where a model trained exclusively on Batch 1 data is applied without retraining.

| **(A) DNA Damage Prediction** |  |  |  |  |  |  |  |  |
| --- | --- | --- | --- | --- | --- | --- | --- | --- |
| Batch | Test Type | Accuracy | Precision | Sensitivity | Specificity | F1-score | AUROC | AUPRC |
| Batch1<br>(8.31%) | From Scratch | 0.99828 | 0.99899 | 0.99789 | 0.99876 | 0.99844 | 0.99997 | 0.99997 |
|  | Direct Test | - | - | - | - | - | - | - |
| Batch2<br>(19.08%) | From Scratch | 0.98950 | 0.99376 | 0.98468 | 0.99410 | 0.98920 | 0.99955 | 0.99955 |
|  | Direct Test | 0.91930 | 0.97225 | 0.85935 | 0.97657 | 0.91232 | 0.98549 | 0.98541 |
| Batch3<br>(35.53%) | From Scratch | 0.98549 | 0.99140 | 0.96997 | 0.99490 | 0.98057 | 0.99912 | 0.99865 |
|  | Direct Test | 0.94079 | 0.92586 | 0.91655 | 0.95548 | 0.92118 | 0.98492 | 0.97928 |
| Batch4<br>(37.08%) | From Scratch | 0.99108 | 0.99488 | 0.98231 | 0.99674 | 0.98856 | 0.99967 | 0.99951 |
|  | Direct Test | 0.94728 | 0.96998 | 0.89331 | 0.98214 | 0.93007 | 0.99094 | 0.98734 |
| **(B) Damage Position Localization** |  |  |  |  |  |  |  |  |
| Batch | Test Type | MAE |  | RMSE |  | Hit@.05 |  |  |
| Batch1<br>(8.31%) | From Scratch | 0.00439 |  | 0.00585 |  | 0.99993 |  |  |
|  | Direct Test | - |  | - |  | - |  |  |
| Batch2<br>(19.08%) | From Scratch | 0.00487 |  | 0.00678 |  | 0.99965 |  |  |
|  | Direct Test | 0.00701 |  | 0.00939 |  | 0.99896 |  |  |
| Batch3<br>(35.53%) | From Scratch | 0.00553 |  | 0.00757 |  | 0.99959 |  |  |
|  | Direct Test | 0.00805 |  | 0.01081 |  | 0.99775 |  |  |
| Batch4<br>(37.08%) | From Scratch | 0.00481 |  | 0.00684 |  | 0.99968 |  |  |
|  | Direct Test | 0.00737 |  | 0.01001 |  | 0.99851 |  |  |

**Table 2:** Dataset statistics for LesionBERT and DamageFormer. (A) Summary of datasets used for damage-aware language model pretraining and downstream finetuning of LesionBERT. (B) Summary of datasets used for multimodal DNA damage detection and localization identification with DamageFormer.

| **(A) Dataset for LesionBERT** |  |  |  |  |  |
| --- | --- | --- | --- | --- | --- |
|  | Split | Total samples | Damaged | Undamaged | Damage types |
| Damage-aware pretraining | Train | 190,056 | 190,056 | 0 | O, S, M, L |
|  | Validation | 10,003 | 10,003 | 0 | O, S, M, L |
|  | Total | 200,059 | 200,059 | 0 | O, S, M, L |
| Damage-aware finetuning | Train | 22,016 | 11,008 | 11,008 | O, S, M, L |
|  | Validation | 7,338 | 3,669 | 3,669 | O, S, M, L |
|  | Test | 7,340 | 3,670 | 3,670 | O, S, M, L |
|  | Total | 36,694 | 18,347 | 18,347 | O, S, M, L |
| **(B) Dataset for DamageFormer** |  |  |  |  |  |
| Multimodal learning:<br>batch 1<br>(ratio: 8.31%) | Train | 1,254,052 | 692,005 | 562,047 | O |
|  | Validation | 179,150 | 98,857 | 80,293 | O |
|  | Test | 358,301 | 197,716 | 160,585 | O |
|  | Total | 1,791,503 | 988,578 | 802,925 | O |
| Multimodal learning:<br>batch 2<br>(ratio: 19.08%) | Train | 2,877,793 | 1,405,877 | 1,471,916 | O |
|  | Validation | 411,113 | 200,839 | 210,274 | O |
|  | Test | 822,228 | 401,680 | 420,548 | O |
|  | Total | 4,111,134 | 2,008,396 | 2,102,738 | O |
| Multimodal learning:<br>batch 3<br>(ratio: 35.53%) | Train | 5,358,682 | 2,023,111 | 3,335,571 | O |
|  | Validation | 765,526 | 289,016 | 476,510 | O |
|  | Test | 1,531,053 | 578,032 | 953,021 | O |
|  | Total | 7,655,261 | 2,890,159 | 4,765,102 | O |
| Multimodal learning:<br>batch 4<br>(ratio: 37.08%) | Train | 5,588,286 | 2,192,895 | 3,395,391 | O |
|  | Validation | 798,327 | 313,271 | 485,056 | O |
|  | Test | 1,596,654 | 626,542 | 970,112 | O |
|  | Total | 7,983,267 | 3,132,708 | 4,850,559 | O |

## Discussion

DNA damage is not uniformly distributed across the genome but emerges from the interplay between chemical reactivity, nucleotide context, chromatin organization, and repair dynamics^59– 61^. Nanopore sequencing provides a unique opportunity to detect such damage directly through lesion-induced perturbations of ionic current^23,62,63^. However, these perturbations are embedded within k-mer-dependent electrical signals and are influenced by neighboring bases within the nanopore sensing window^28,64^, making lesion detection intrinsically context dependent. Our findings demonstrate that resolving this ambiguity requires explicit integration of biophysical signal measurements with the intrinsic sequence environment in which lesions occur. In this study, we developed DamageFormer, a damage-aware multimodal deep learning framework for DNA lesion identification based on nanopore sequencing that integrates nucleotide sequence context with raw nanopore electrical signals to enable accurate detection and precise localization of DNA lesions. Through systematic benchmarking, ablation analyses, and cross-dataset validation, we demonstrate that joint modeling of sequence and signal substantially enhances both detection robustness and localization precision compared to unimodal approaches. Beyond performance improvements, our results show that multimodal representation learning captures biologically coherent relationships among lesion chemistry, electrophysiological perturbations, and nucleotide context, providing a scalable and interpretable framework for high-resolution DNA damage mapping.

A central challenge in genome-wide DNA damage profiling is the extremely low prevalence of lesion events relative to the vast number of genomic bases^65–67^. For example, steady-state levels of oxidative lesions, such as 8-oxo-dG, are estimated at approximately 0.5-2.5 lesions per million base pairs under physiological conditions, making target events rare against a background of normal bases. In practical applications, even a small false positive rate can yield thousands to millions of erroneous calls when extrapolated across large genomic regions. This accumulation of errors not only increases downstream validation burden but also obscures true biological signals, particularly in studies aiming to link damage patterns to mutational processes, repair pathway engagement, or disease phenotypes^59,68,69^. Consequently, performance improvement that may appear numerically modest in standard metrics (e.g., small gains in F1-score or incremental reductions in MAE) can correspond to substantial decreases in false discoveries at the genome scale. From this perspective, achieving near-optimal performance is not a marginal optimization but a fundamental requirement for reliable biological interpretation and scalable deployment. A key mechanistic advance of this work lies in the construction of the genomic foundation model LesionBERT that enables damage-aware pretraining of biological sequences. Oxidative lesions such as 8-oxo-7,8-dihydro-2′-deoxyguanosine (8-oxo-dG) alter hydrogen bonding geometry and promote Hoogsteen base pairing, increasing the likelihood of *G* → *T* transversions during replication^70–72^. In contrast, O6-alkylguanine adducts differ in steric bulk and electronic structure, influencing polymerase progression and mutagenic outcomes in lesion-specific ways^7,73,74^. These biochemical differences are reflected not only in signal perturbations measurable by nanopore sequencing^62,63^ but also in sequence-context dependencies and repair pathway engagement, including base excision repair and O6-methylguanine-DNA methyltransferase (MGMT)-mediated reversal^75,76^. By extending the nucleotide vocabulary to encode lesion categories and applying asymmetric masking during pretraining, LesionBERT is explicitly capable of learning contextual dependencies surrounding modified guanines. This strategy parallels recent advances in biological foundation models, where domain-adapted pretraining enhances downstream specialization^40,41^. The consistent improvement of detection performance observed when replacing generic nucleotide embeddings with damage-aware representations indicate that contextual sequence features encode biologically meaningful constraints that stabilize the interpretation of noisy electrical signals.

The multimodal integration framework further reflects the dual nature of lesion detection. Nanopore signals directly capture electrophysiological consequences of chemical modification, including altered current amplitude and dwell time distributions^23,64^. Yet these signals propagate across multiple adjacent bases within the pore, producing spatial diffusion of perturbations^28^. Nucleotide sequence context provides complementary structural and biochemical information that constrains lesion plausibility and help distinguish genuine damage-induced deviations from canonical base-dependent signal variation. The adaptive fusion mechanism demonstrates that the relative importance of sequence and signal varies across molecules, consistent with the biological heterogeneity of lesion manifestation in **Fig. 5C**. Our systematic loss-weighting analysis further highlights that moderate cross-modal contrastive alignment enhances representational coherence between sequence and signal embeddings, consistent with advances in multimodal and self-supervised learning^47,48^. However, excessive alignment pressure degrades detection and localization accuracy, suggesting that over-constraining modality agreement can suppress lesion-specific discriminative features in **Fig. 6**. Likewise, increasing positional localization weight improves boundary precision only up to an optimal threshold, beyond which detection stability declines. This trade-off mirrors the physical constraints of nanopore sensing, in which ionic current reflects a sliding k-mer window rather than a single nucleotide^28,64^. Visualization of the learned embedding space in **Fig. 4E** provides additional insight into model behavior. We further assessed the effect of the hard-negative-aware contrastive loss (Additional file 1: **Fig. S2**). Relative to alternative contrastive objectives (Additional file 2) and the no-contrastive baseline, hard-negative mining produced more discriminative cross-modal representations and yielded consistent improvements in both detection accuracy and localization precision. Interpretability analyses further support biological plausibility in **Fig. 7**, where in damaged samples, both gradient-based attribution and in-silico mutagenesis localize predictive evidence to narrow sequence-signal regions, while undamaged samples exhibit diffuse attribution patterns. Evaluation on the independent 8-oxo-dG dataset generated under distinct experimental settings demonstrates scalable generalizability in **Table 1**, indicating that the model captures fundamental chemical and contextual properties of guanine oxidation rather than dataset-specific artifacts.

Despite these strengths, several limitations warrant consideration. First, the model was trained primarily on synthetic oligonucleotide data containing site-specific DNA adduct insertion. Although this design enables precise ground-truth labeling, it may not fully capture the complexity of damage in native genomic contexts, where chromatin structure, DNA-protein interaction, and epigenetic modifications influence both lesion formation and nanopore signal characteristics. Second, the proposed framework operates on fixed-length sequence and signal windows, restricting its ability to capture long-range genomic dependencies that may influence damage susceptibility or repair. Emerging architectures designed for long biological sequences^38,77^ offer promising directions for extending contextual range in future iterations. Third, while synthetic constructs are necessary due to the scarcity of gold-standard lesion annotations in genomic DNA^24,25,27^, broader validation in native genomic contexts, encompassing the chromatin structure^78^ and epigenetic modifications^79^, will be essential to confirm generalizability under biologically realistic conditions. Finally, interpretability analyses can provide qualitative insights into modality reliance and feature importance. However, they remain inherently model-dependent and do not substitute direct biological experimental validation. Integrating orthogonal wet-lab evidence or perturbation-based validation will be important for translating computational predictions into mechanistic biological insight.

## Conclusions

We developed DamageFormer, a multimodal deep learning framework that integrates DNA sequence context with raw nanopore signals to enable accurate detection and localization of DNA lesions. By introducing LeisonBERT, a damage-aware genomic foundation language model, and coupling it with adaptive gated cross-modal fusion, our model achieves superior performance in both lesion detection and positional localization compared with existing state-of-the-art baseline methods. Systematic ablation and embedding analyses demonstrate that damage-aware pretraining, balanced auxiliary supervision, and multimodal alignment are all critical for robust rare-event genomic inference. Beyond performance improvement, DamageFormer provides biologically coherent representations that capture structured relationships among lesion chemistry, electrophysiological perturbations, and nucleotide context. The model exposes class structure directly within embedding geometry, maintains stable optimization across hyperparameter regimes, and enhances interpretability through spatially localized attribution signals. In summary, this work offers a scalable and transferable framework for high-resolution DNA damage mapping, advancing the incorporation of chemical DNA information into next-generation genomic analysis and opening new opportunities for studying mutational processes, genome instability, and disease-associated damage landscapes.

## Methods

### Dataset construction

#### Pretraining and Finetuning Dataset

For damage-aware pretraining and downstream finetuning, we used nanopore sequencing datasets from multiple published sources^27,80–86^ that systematically characterize DNA damage signatures across chemically distinct guanine adducts. Synthetic plasmid DNA was exposed to oxidizing and alkylating agents to generate eleven well-defined lesions, including the oxidative lesion 8-oxo-7,8-dihydro-2′-deoxyguanosine (8-oxo-dG) and a diverse set of O6-alkylguanine adducts spanning varying steric sizes and chemical properties. Rather than modeling each adduct separately, lesions were grouped into four biologically meaningful categories represented by discrete tokens: oxidative (O), small alkyl (S), medium alkyl (M), and large bulky alkyl (L). This design was motivated by biochemical relevance and the nanopore signal. Oxidative lesions such as 8-oxo-dG induce characteristic *G* → *T* transversions through Hoogsteen pairing^5^, whereas O6-alkylguanine adducts form a steric continuum ranging from small substitutions causing minimal helix disruption to bulky groups that strongly perturb DNA geometry and polymerase activity^81,83,84^. Importantly, nanopore studies show that ionic current perturbations correlate more strongly with adduct size class than precise chemical identity, while structurally similar adducts often produce overlapping signal signatures^27^. Grouping lesions into four classes, therefore, preserves major axes of biological variation while avoiding sparse vocabularies that hinder stable embedding learning^8,87,88^.

Damaged plasmids were sequenced using Oxford Nanopore Technologies (ONT) MinION devices, yielding both raw current traces and basecalled reads. Reads were aligned to reference plasmid sequences, and damage positions were determined from known modification sites. Paired sequence-signal samples were extracted by defining variable-length sequence windows centered on lesion sites together with corresponding nanopore signal segments. Undamaged controls were generated from matched genomic regions lacking modifications to maintain comparable sequence context and signal-length distributions. Damage positions were normalized to the unit interval as:

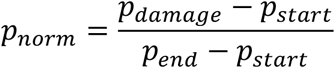

where *p*_*damage*_ is the absolute damage position and [*p*_*start*_, *p*_*end*_] defines the sequence window boundaries. We ended up with 218,406 damaged and 1,041,420 undamaged samples (**Table 2A**). For damage-aware language model pretraining, only damaged samples were used, yielding 200,059 samples split into training and validation sets (95:5). The pretraining distribution comprised 41.2% small alkyl, 34.2% medium alkyl, 15.8% large bulky, and 8.8% oxidative lesions. For supervised finetuning and evaluation, balanced datasets were created by randomly subsampling undamaged reads at a 1:1 ratio. Data were split into training, validation, and test sets at a 6:2:2 ratio, producing 22,016 training, 7,338 validation, and 7,340 test samples.

#### Multimodal DNA Damage Dataset

To evaluate the generalization of DamageFormer, we used an independent nanopore sequencing dataset profiling 8-oxo-dG lesions^26^. This dataset comprises four independently constructed batches of synthetic oligonucleotide samples containing site-specific 8-oxo-dG modifications sequenced on Oxford Nanopore Technologies (ONT) platforms. The batches differ in size and experimental conditions, enabling assessment of cross-batch robustness and scalability. The proportions of the total dataset represented by Batch 1-4 are 8.31%,19.08%, 35.53%, and 37.08%, respectively. Among these, Batch 1 served as the primary dataset for model development. Raw nanopore signal traces and corresponding basecalled reads from Batch 1 were processed using the same preprocessing pipeline as the primary dataset to extract paired sequence-signal windows centered on annotated lesion sites or matched undamaged control positions. Batch 1 was divided into training (70%), validation (10%), and test (20%) subsets using stratified sampling to preserve class balance. Only the held-out test subset from Batch 1 was used for primary performance evaluation. The remaining three batches (Batches 2-4) were treated as independent external validation datasets. These batches were processed based on the identical preprocessing pipeline but were not used during training. The performance on Batches 2-4 reflects the model’s ability to generalize across independent experimental replicates and diverse data distributions. Detailed statistics of all dataset splits are provided in **Table 2B**.

### The creation of LesionBERT

We first developed LesionBERT, a damage-aware foundation genomic language model, through a two-stage training strategy that was designed to encode damage-aware sequence representations for damage detection. In the first stage, we performed continued pretraining of DNABERT-2 using a damage-aware masked language modeling objective with an extended vocabulary and asymmetric masking to emphasize lesion contexts. In the second stage, the pretrained model was finetuned for DNA damage detection using Low-Rank Adaptation (LoRA), enabling parameter-efficient task adaptation while preserving the learned damage-sensitive representations.

#### Damage-Aware Language Model Pretraining

(1) Vocabulary extension: To enable damage-aware representation learning, we extended the DNABERT-2 vocabulary by introducing four damage-specific tokens corresponding to chemically grouped lesion categories: O (oxidative damage; 8-oxo-dG), S (small alkyl adducts: O6-methyl-dG, O6-ethyl-dG, O6-hydroxyethyl-dG), M (medium alkyl adducts: O6-n-propyl-dG, O6-isopropyl-dG, O6-n-butyl-dG, O6-isobutyl-dG, O6-s-butyl-dG), and L (large/bulky adducts: O6-neopentyl-dG, O6-benzyl-dG). Although eleven distinct guanine adducts were experimentally profiled, we introduced four specialized tokens instead of assigning one token per lesion type. This is due to the resolution limits of nanopore signal measurement and principled modeling consideration. Nanopore current perturbations primarily reflect steric class and local structural distortion within the pore sensing window, and structurally similar adducts often produce overlapping signal profiles that are not reliably separable at the individual substituent level. Assigning a distinct token to each lesion would exceed the effective resolution of the signal modality. In addition, several adduct types occur at relatively low frequencies; allocating individual tokens to each lesion would result in sparse supervision and unstable embedding learning during masked language modeling. Instead, grouping lesions into oxidative, small, medium, and bulky classes captures the primary biochemical axes of variation, oxidation versus alkylation and increasing steric bulk, while ensuring sufficient token frequency for stable and robust language model optimization. These four tokens were added as special vocabulary items, increasing the vocabulary size from 4,096 to 4,100. The input embedding matrix **E** and the masked language modeling (MLM) output projection were resized accordingly. Damage token embeddings were initialized from the pretrained guanine embedding with small Gaussian perturbations:

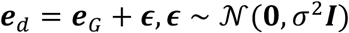

where **e**_*G*_ denotes the guanine embedding, *σ* is the standard deviation controlling noise magnitude (*σ* = 0.01), and ***I*** is the identity matrix. This initialization reflects the biochemical origin of the lesions as guanine modifications while allowing the model to differentiate damage categories during continued pretraining. The small variance ensures stability by keeping damage embeddings close to canonical guanine in the representation space. (2) Model architecture: LesionBERT was implemented as a MLM consisting of the DNABERT-2 encoder and a customized MLM prediction head. The encoder follows the standard DNABERT-2 transformer architecture (12 layers, hidden size 768, 12 attention heads; approximately 117M parameters). The MLM head maps hidden states to vocabulary logits through a two-layer projection:

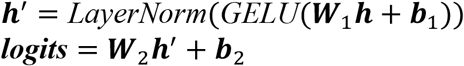

where **h** is the hidden state at each position, **W**_1_, **W**_2_, **b**_1_, **b**_2_ are the learnable parameters, and GELU (Gaussian Error Linear Unit) is a smooth nonlinear activation function. (3) Asymmetric masking strategy: To emphasize learning of rare lesion tokens, we used an asymmetric masking scheme with token-dependent probabilities:

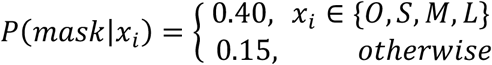

Special tokens ([PAD], [CLS], [SEP]) and padding positions were excluded from masking. Masked tokens followed the standard BERT replacement scheme: 80% replaced with [MASK], 10% replaced with a random token, and 10% left unchanged. The MLM objective minimized cross-entropy loss over masked positions:

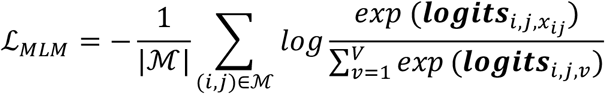

where ℳ denotes the set of masked positions. The elevated masking rate for lesion tokens (40% vs 15%) ensures that these rare events contribute sufficiently to the training signal. (4) Training details: Continued pretraining was performed using the AdamW optimizer with learning rate 1 × 10^−5^, weight decay 0.01, and linear warmup over the first 10% of training steps, followed by linear decay. Training ran for 10,000 steps using batch size 32 per L4 GPU across three GPUs (effective batch size 96), maximum sequence length 512, mixed-precision training (FP16), and gradient clipping with maximum norm 1.0. Model checkpoints were saved every 5,000 steps.

#### Damage-Aware Language Model Pretraining

(1) Model instantiation and task head construction: The pretrained LesionBERT encoder was initialized with the extended vocabulary and pretrained weights. A two-class prediction head was added on top of the pooled sequence representation (corresponding to the [CLS] token embedding). The prediction head consists of a linear projection mapping the 768-dimensional pooled representation to binary logits. All token embeddings were verified to match the extended vocabulary introduced during pretraining. (2) Parameter-efficient adaptation with LoRA: To enable task adaptation while preserving pretrained representations, we applied Low-Rank Adaptation (LoRA)^89^. Instead of updating the full transformer weights, LoRA reformulates weight updates as:

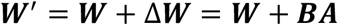

where **W** ∈ ℝ^*d×k*^ denotes the frozen pretrained weight matrix, and **B** ∈ ℝ^*d×r*^ and **A** ∈ ℝ^*r×k*^ are trainable low-rank matrices with rank *r* ≪ min (*d, k*). LoRA adapters were inserted into the query, key, and value projection matrices of all self-attention layers. We used rank *r* = 16, scaling factor *α* = 32, and dropout rate 0.05. Under this configuration, only a small fraction of parameters was updated during finetuning, while the core transformer weights remained frozen. This design improves training stability and reduces overfitting, particularly under limited labeled data. (3) Supervised training objective: The model was trained for the detection of damaged sequences. The primary objective was binary cross-entropy (BCE) loss between predicted logits and ground-truth labels. (4) Optimization settings: Finetuning was performed using the AdamW optimizer with separate learning rate scaling for LoRA parameters and prediction head weights. The pretrained encoder parameters remained frozen except for LoRA adapters. Training employed mixed precision, gradient clipping, and early stopping based on validation performance. Data were split into training, validation, and test subsets as described in the Dataset section.

### The architecture of DamageFormer

To jointly exploit sequence context and raw nanopore signal information, we proposed a multimodal deep learning framework for DNA damage detection that integrates sequence context and electronic signal information in a unified and end-to-end manner (**Figure 1**). The sequence encoder, i.e., LesionBERT, learns contextual patterns in the DNA sequence, while a parallel signal encoder captures local fluctuations in the associated electronic signal indicative of damage. The generated modality-specific embeddings are adaptively combined through a learned fusion gate, which dynamically weights the contributions of sequence and signal features for each sample. The fused representation is then shared by two task-specific heads for damage detection and damage position localization, enabling joint learning of presence and localization. Training is guided by a combination of prediction loss, position localization loss applied to damaged samples, and a contrastive loss that aligns sequence and signal representations, promoting cross-modal consistency.

#### Sequence Encoder

The sequence encoder, called LesionBERT, consists of the finetuned DNABERT-2 model with LoRA adapters merged into the base weights. For an input sequence **x** = (*x*_1_, *x*_2_, …, *x*_*n*_), the encoder produces contextualized representations:

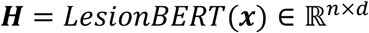

where *d* = 768 is the hidden dimension. We extract the [CLS] token representation as the sequence-level embedding ***h***_*seq*_ = ***H***[0], which is then projected to the fusion dimension:

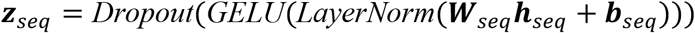

where **W**_*seq*_ and **b**_*seq*_ are the weights and bias to be learned for the projection layer.

#### Signal Encoder

The signal encoder processes raw nanopore current measurements **s** = (*s*_1_, *s*_2_, …, *s*_*T*_) ∈ ℝ^*T*^ through a CNN-BiLSTM architecture with the residual connection designed to capture both local signal patterns and long-range temporal dependencies. Nanopore signals exhibit characteristic current fluctuations at damage sites, but these perturbations are embedded within noisy baseline currents that vary with sequencing conditions. The encoder, therefore, combines convolutional layers to extract local signal motifs with recurrent layers to model temporal context. (1) Convolutional feature extraction: Two 1D convolutional layers with residual connection extract hierarchical signal features:

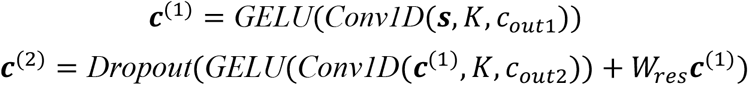

where the kernel size is K, and the output dimensions are *c*_*out*1_ and *c*_*out*2_, **W**_*res*_ is a convolution layer that projects the first-layer features to match the channel dimension. We also added a residual connection layer, which connects the outputs of the convolution layers:

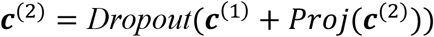

where *Proj* is the same as the mentioned projection layer. This residual connection facilitates gradient flow and allows the second layer to learn refinements over the initial features rather than complete transformations, which stabilizes training for the relatively shallow signal encoder. In our experiments, K=7, *c*_*out*1_ = 3_2_, *c*_*out*2_ = 64, and **W**_*res*_ is a 1×1 convolution kernel. Both layers use the same padding to preserve temporal resolution. (2) Recurrent sequence modeling: A single-layer bidirectional LSTM per direction processes the convolutional features:

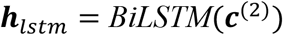

The bidirectional architecture enables the model to incorporate both upstream and downstream signal context when characterizing each position, which is important because damage-induced current perturbations can extend beyond the immediate lesion site. The signal representation is obtained by mean pooling across the temporal dimension:

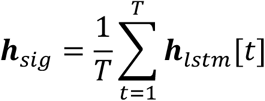

(3) Projection: The pooled representation is projected to the fusion dimension through a linear layer with layer normalization and GELU activation:

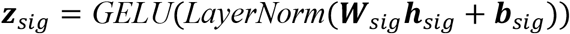

where **W**_*sig*_ and **b**_*sig*_ are learnable parameters for the signal projection.

#### Multimodal Fusion

To adaptively combine sequence and signal representations, we employ a learned gating mechanism that dynamically determines the relative contribution of each modality for each input sample. The gate network takes the concatenated embeddings as input and produces fusion weights through a two-layer multilayer perceptron (MLP):

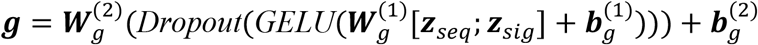

where 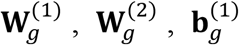 and 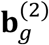 are learnable parameters. Temperature-scaled SoftMax normalization yields fusion weights for the sequence and signal modals **w** = *softmax*(***g***/*τ*), where *τ* is a temperature parameter controlling the sharpness of the weight distribution^90^. The fused representation is computed as the weighted combination:

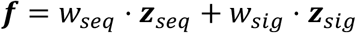

This fusion method enables the model to emphasize sequence context when signal quality is poor or to prioritize signal features when electrical perturbations are pronounced, providing an interpretable mechanism for understanding modality contributions across different damage types and experimental settings.

#### Prediction Heads

The fused representation **f** is passed to two task-specific prediction heads that jointly address damage detection and localization. (1) Prediction head: A 2-layer MLP produces binary prediction logits for damage detection:

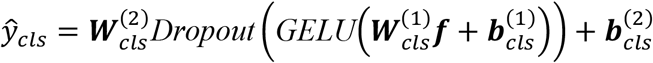

where 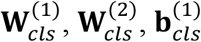 and 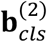 are learnable parameters. The intermediate bottleneck dimension reduces overfitting while retaining sufficient capacity to distinguish damaged from undamaged DNA samples. (2) Position localization head: A 2-layer MLP with sigmoid activation identifies the normalized damage position within the sequence window:

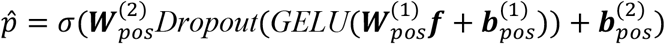

where *σ*(⋅) is the sigmoid function constraining predictions to 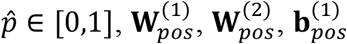 and 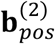 are learnable parameters. The position head shares the same architecture as the prediction head but is trained only on damaged samples with valid position annotations, treating localization as an auxiliary task that provides additional supervision for learning damage-sensitive representations.

### Training objective

The model is trained with a multi-task objective that combines prediction, position localization, and contrastive alignment losses. Each component addresses a distinct challenge in damage detection, i.e., class imbalance, precise localization, and multimodal representation learning. This joint objective enables stable optimization and balanced performance across detection and localization tasks. The details can be found as follows.

#### Multi-Task Loss Function

The total training loss combines three complementary objectives formulated below:

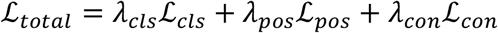

Here *λ*_*cls*_, *λ*_*pos*_, and *λ*_*con*_ are the coefficients to balance the contributions of each part. The prediction loss receives the highest weight as the primary objective, while position localization and contrastive losses provide auxiliary supervision that improves learned representations without dominating the training dynamics.

#### Focal Loss for Prediction

Damaged samples typically constitute a minority of sequencing reads, creating a class imbalance that can bias standard cross-entropy toward predicting the majority class. We address this with the focal loss^91^, which down-weights well-classified examples to focus learning on hard samples:

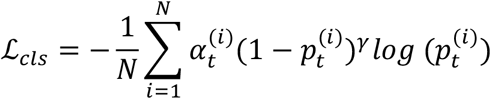

where *N* denotes the number of training samples. The term 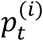 represents the model’s estimated probability for the ground-truth class:

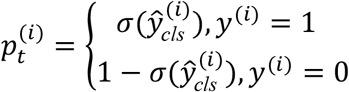

and 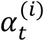 is a class-balancing factor defined as:

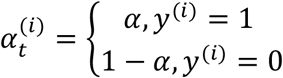

The parameter *γ* modulates the rate at which easy examples are downweighed (by default as 2.0). When a sample is well-classified (*p*_*t*_ → 1), the factor (1 − *p*_*t*_)^*γ*^ approaches zero, reducing its contribution to the loss. This allows the model to concentrate on ambiguous or misclassified examples where additional learning is most beneficial.

#### Smooth L1 Loss for Position Localization

For damage position localization, we employed the Smooth L1 (Huber) loss^92^, applied only to damaged samples with valid position annotations. Specifically, the loss is defined over the set 𝒫 = {*i* | *y*^(*i*)^ = 1 ∧ *p*^(*i*)^ ≥ 0}, and is computed as:

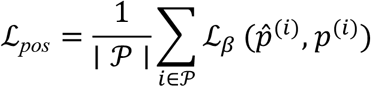

where 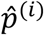 and *p*^(*i*)^ denote the predicted and ground-truth damage positions, respectively. The Smooth L1 loss is given by:

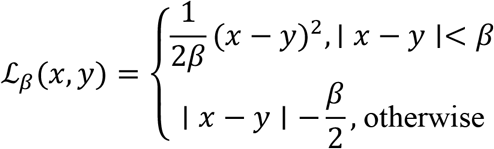

where β is the transition parameter controlling the boundary between quadratic and linear regimes. In our experiments, β was set to 0.05, corresponding to 5% of the normalized sequence window length. The quadratic regime near zero provides smooth gradients that facilitate precise positioning when predictions are close to targets, while the linear regime for larger errors prevents outliers from producing excessively large gradients that could destabilize training.

#### Hard-Negative-Aware Contrastive Loss

To learn aligned multimodal representations that are both discriminative and robust, we employed a contrastive loss with hard negative mining. Unlike the standard InfoNCE loss function^93^ that treats all negatives equally, we explicitly identify the most challenging negative samples, i.e., those undamaged samples that are highly similar to damaged cases. Given a batch of *B* samples with L2-normalized sequence embeddings 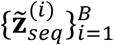 and signal embeddings 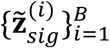, we first compute the pairwise similarity matrix:

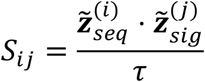

where *τ* is the temperature parameter. For each sample *i*, the positive similarity was defined as the similarity between its matched sequence and signal embeddings: 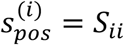. To construct hard negatives, we defined a label-disagreement mask **D**, where 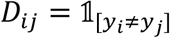 indicates whether samples *i* and *j* have different damage labels. For each sample *i*, the hardest negative is selected as the label-mismatched pair that exhibits the highest similarity:

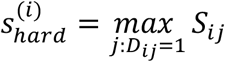

This corresponds to the most confusable negative example, i.e., the mismatched sequence-signal pair whose representation is most similar to the anchor sample. The loss function incorporates the hard negative with a weighting factor *λ*_*hard*_:

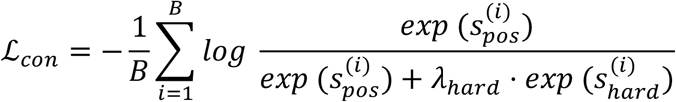

### Model robustness and interpretability

To assess the robustness and scalability of DamageFormer, we conducted a controlled multi-batch study using four independently constructed datasets that differ substantially in size. The performance was evaluated under two regimes: (1) training and testing from scratch within each batch to examine stability across dataset scales, and (2) directly applying a model trained on a single dataset to the remaining batches without retraining to test cross-dataset generalization. This design isolates the model’s ability to transfer damage-aware representations learned from limited data to larger and distributionally distinct datasets. To elucidate the underlying mechanisms driving these predictions, we performed complementary interpretability analyses at both dataset-wide and individual-sample levels. At the dataset level, we quantified modality reliance by analyzing the distribution of fusion gate weights and examining their relationship to prediction confidence. To test whether fusion reflects functional dependence rather than passive correlation, we conducted cross-modal swap experiments in which either sequence or signal inputs were permuted across samples and the resulting changes in predicted probabilities were measured. At the sample level, we applied integrated gradients to quantify feature importance for both modalities. For signal attribution, gradients were accumulated along a linear interpolation between a zero baseline and the observed nanopore current trace with respect to the prediction output. For sequence attribution, integrated gradients were computed over input token embeddings of the sequence encoder while holding the signal branch fixed, isolating nucleotide-level contributions. Finally, we performed in-silico mutagenesis by systematically substituting nucleotides within a defined sequence window and measuring the corresponding changes in predicted probability of DNA damage. Collectively, these analyses provide a multi-scale assessment of model robustness, cross-modal interaction, and damage localization evidence.

### Benchmark comparison

To comprehensively evaluate the performance of DamageFormer, we compared it against a diverse set of baseline approaches spanning nanopore-specific deep learning models, MLP-based deep learning algorithms, and traditional machine learning classifiers. All baselines were trained and evaluated under identical procedures and protocols in the same setting to ensure fair comparison.

#### (1) Nanopore-specific deep learning baselines

We benchmarked against several representative nanopore-based frameworks, including DeepSignal^24^, NanoCon^43^, Oxo^26^, DeepMod^56^, and Tombo^94^. DeepSignal is a multimodal nanopore modification detection framework that combines convolutional neural networks and bidirectional LSTM modules to capture local signal patterns and sequential dependencies. NanoCon employs a hybrid neural architecture incorporating contrastive learning to align sequence-derived and signal-derived representations. Oxo is a deep learning framework that integrates convolutional encoders for local signal feature extraction with recurrent modules to capture temporal dependencies, enabling joint analysis of signal and sequence information. DeepMod is a signal-only modification detection model based on bidirectional LSTM networks that model temporal dependencies in raw nanopore current traces. Tombo is a nanopore signal analysis framework that combines neural signal feature extraction with contextual aggregation strategies inspired by statistical deviation scoring to detect modification-associated signal shifts.

#### (2) MLP-based deep learning baselines

We designed three MLP variants to evaluate the impact of model depth and encoder adaptation on lesion prediction. Light MLP is a lightweight sequence-only baseline that applies a single linear projection (or shallow MLP) to pretrained sequence embeddings extracted from a genomic language model. Deep MLP extends this configuration by incorporating multiple fully connected layers with nonlinear activations, normalization, and dropout regularization to increase representational capacity while still operating on fixed embeddings. Finetuned MLP further allows end-to-end optimization of both the finetuned sequence encoder and the MLP prediction head. By updating encoder parameters during training, this baseline evaluates whether task-specific adaptation of sequence representations can improve lesion prediction compared with frozen-encoder models.

#### (3) Traditional machine learning baselines

We utilized several widely used classic machine learning classifiers, including XGBoost, random forest (RF), linear regression (LR), and support vector machine (SVM). For these models, input features consisted of pretrained sequence embeddings and aggregated signal-derived features to ensure comparability with neural baselines. All classifiers were implemented using standard library implementations and trained with default hyperparameter settings, without extensive tuning, to establish transparent and reproducible baseline performance.

### Evaluation metrics

#### Metrics for DNA lesion prediction

For DNA damage detection, we evaluated model performance using threshold-dependent metrics, including precision, sensitivity, specificity, and F1-score. To assess discriminative ability independent of a fixed prediction threshold, we additionally reported the area under the receiver operating characteristic curve (AUROC) and the area under the precision-recall curve (AUPRC), which summarize performance across all possible decision thresholds.

#### Metrics for DNA position prediction

For damage localization evaluation, we assessed the performance on the subset of correctly classified positive samples, denoted as 𝒫_*coorect*_:

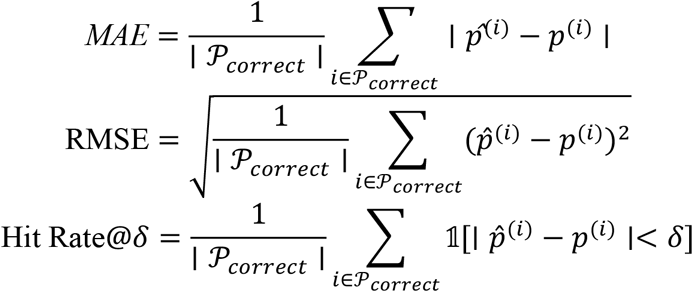

We reported Hit Rate@0.05, corresponding to predictions within 5% of the true normalized position. For 128-nucleotide sequence windows, this threshold corresponds to approximately ±6 nucleotides, providing a practical measure of sub-read localization accuracy.

## Supporting information

Contrastive loss functions

## Author information

### Contributions

R.Y. and Q.M. conceived this project. Q.Y. designed the code and performed experiments. L.L. prepared and analyzed the data. Q.Y. wrote the manuscript. R.Y., and Q.M. revised the manuscript.

### Data availability

The data and codes underlying this article are publicly available through GitHub at https://github.com/UF-HOBI-Yin-Lab/DamageFormer.

## Ethics declarations

### Ethics approval and consent to participate

Not applicable.

### Consent for publication

Not applicable.

## Supplementary Information

Additional file1: Figures S1-S2: Supplementary figures S1 and S2. Additional file2: Contrastive loss functions

