## Supplementary material for "DamageFormer: a damage-aware multimodal deep learning framework for DNA lesion identification from nanopore sequencing": Figures S1-S2

### Additional file 1: Figures S1-S2

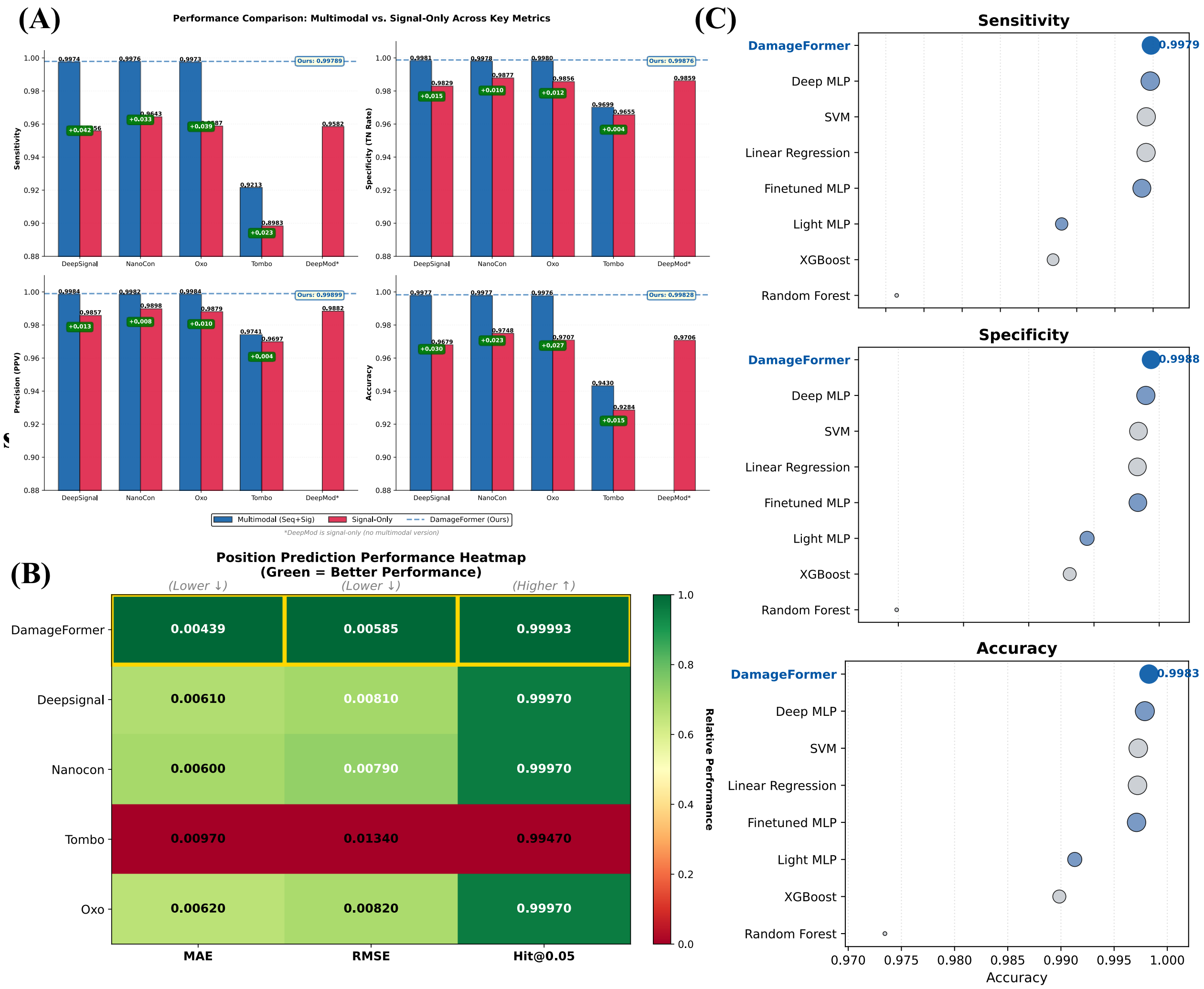


**Fig. S1**: The performance comparison of DamageFormer and baseline models. (A) Multimodal models consistently achieved better performance compared to signal data-based models in term of sensitivity, specificity, precision, and accuracy. (B) DamageFormer got the best performance on damage position prediction measured by mean absolute error (MAE), root mean square error (RMSE), and Hit@0.05 accuracy. (C) DamageFormer ranked first among deep learning baselines, and traditional machine learning methods for DNA damage prediction across sensitivity, specificity, and accuracy.

### Results

#### Multimodal models outperform signal data-based models

To quantify the contribution of multimodal integration, we further demonstrated that multimodal models consistently outperform their signal-only counterparts compared to signal-only learning in terms of sensitivity, specificity, precision, and accuracy across different models in **Fig. S1A**. Incorporating sequence context alongside nanopore signal information yielded a systematic increase in detection reliability. These improvements were most pronounced for sensitivity and precision, indicating enhanced ability to detect true lesion events while reducing false positives. Specificity and overall accuracy showed the same directional trend, confirming that multimodal integration improves global classification stability rather than optimizing a single metric. Importantly, the separation between multimodal and signal-only models persisted in a nearoptimal performance regime, indicating that these improvements reflected genuine modeling advantages rather than random fluctuations.

**DamageFormer outperforms baselines in lesion position prediction**

**Fig. S1B** compared lesion localization accuracy across competing methods using mean absolute error (MAE), root mean square error (RMSE), and tolerance-based Hit@0.05. DamageFormer achieved the lowest error across both MAE and RMSE while simultaneously attaining the highest tolerance-based accuracy, establishing a consistent performance advantage across all localization criteria. The heatmap highlighted a clear separation between DamageFormer and the baseline models, indicating reduced positional variance and more stable boundary estimation around damaged bases. Competing approaches showed higher error and greater variation across metrics, suggesting less reliable alignment between signal perturbations and genomic coordinates. In contrast, DamageFormer maintains stability across error-based and tolerance-based measures.

**DamageFormer consistently outperforms classical ML and generic DL models**

Across sensitivity, specificity, and accuracy, DamageFormer consistently shows the upper performance boundary relative to both classical machine learning and generic deep learning baselines (**Fig. S1C**). Traditional classifiers such as Random Forest, XGBoost, SVM, and linear regression exhibit noticeably lower performance, suggesting limited capacity to capture complex interactions between nanopore signal dynamics and sequence context. Generic neural architectures improve upon these methods but remain separated from DamageFormer by a persistent margin across all metrics. Notably, the performance advantage is consistent across metrics rather than confined to a single measure. DamageFormer achieves the highest sensitivity, specificity, and accuracy, indicating balanced discrimination. This pattern suggests that the observed improvements stem from domain-specific multimodal representation learning instead of increased model capacity alone.

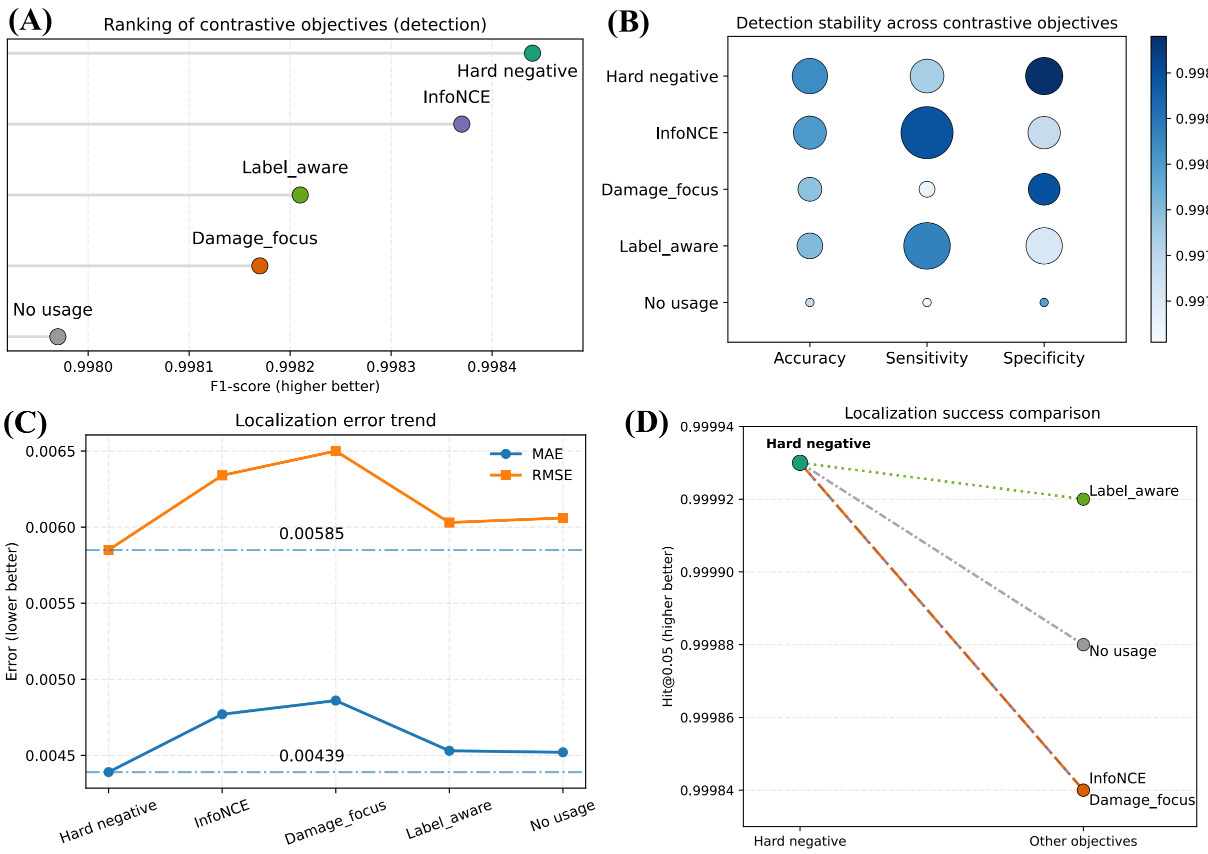


**Fig. S2**: Systematic comparison of contrastive learning objectives for multimodal DNA damage prediction and localization. (A) Ranking of contrastive objectives based on detection performance measured by F1-score. Each point represents a training objective, ordered from lowest to highest performance, highlighting the superiority of hard-negative contrastive learning. (B) Bubble heatmap summarizing detection stability across accuracy, sensitivity, and specificity. Color intensity encodes absolute performance, while bubble size reflects improvement relative to the no-contrastive baseline. (C) Localization error trend across contrastive objectives. Mean absolute error (MAE) and root mean squared error (RMSE) quantify regression precision, with lower values indicating improved positional accuracy. (D) Comparison of localization success measured by Hit@0.05. Lines connect each objective to the hard-negative baseline, illustrating relative changes in high-precision localization.

#### Hard-negative-aware contrastive loss function beats other types of loss functions

We evaluated several contrastive learning strategies, including standard InfoNCE, label-aware contrastive loss, and damage-focused contrastive loss in **Fig. S2**. The hard-negative-aware contrastive loss consistently delivered the best performance, particularly in improving discrimination between damaged and undamaged samples with similar signal or sequence patterns. By explicitly emphasizing challenging negative pairs, this loss function encourages more discriminative cross-modal embeddings and leads to better generalization in both detection and localization tasks.

**Fig. S2A** compares damage detection performance across contrastive objectives using F1-score ranking. Hard-negative contrastive learning achieved the highest score, outperforming InfoNCE, label-aware, damage-focused, and the no-contrastive baseline. Although differences were numerically modest, the consistent ordering indicated that explicitly mining difficult cross-modal negatives provides stronger representation separation than generic alignment strategies. Label-aware and damage-focused objectives yielded moderate improvements over the baseline, suggesting that incorporating task structure helps but is insufficient without emphasizing hard negatives. Overall, the results show that contrastive design directly influences multimodal feature discriminability. **Fig. S2B** assesses detection stability across metrics using a bubble heatmap that encodes both absolute performance and improvement magnitude. Hard-negative learning produced the most balanced gains across accuracy, sensitivity, and specificity. In contrast, InfoNCE and other variants introduced metric trade-offs, improving certain axes while degrading others. The no-contrastive setting consistently underperformed, confirming that contrastive supervision enhances multimodal alignment. Importantly, hard-negative mining improved all metrics coherently rather than producing isolated gains, indicating stronger decision boundary separation.

**Fig. S2C** examines localization error. Hard-negative contrastive learning achieved the lowest MAE and RMSE, demonstrating superior positional precision. Other objectives increased regression error to varying degrees, with InfoNCE and damage-focused losses showing the largest degradation. Label-aware learning partially mitigated this effect but did not match hard-negative performance. These findings indicate that contrastive objective choice directly affects localization robustness. **Fig. S2D** shows that localization accuracy (Hit@0.05) follows the same trend: hard-negative learning establishes the highest precision, while alternative objectives yield incremental declines. These results demonstrate that selectively emphasizing hard negatives sharpens embedding separation and translates into consistent improvements in both detection and localization performance.
