## Supplementary material for "DamageFormer: a damage-aware multimodal deep learning framework for DNA lesion identification from nanopore sequencing": Contrastive loss functions

### Additional file 2: Contrastive loss functions

#### Methods

##### Comparative contrastive loss functions

To encourage alignment between sequence-derived and signal-derived representations, we investigated several contrastive learning objectives that have been widely used in multimodal and self-supervised representation learning. Contrastive learning aims to pull semantically related embeddings closer while pushing unrelated embeddings apart, thereby structuring the embedding space according to task-relevant relationships.

*InfoNCE Loss*. As a baseline contrastive objective, we implemented the InfoNCE loss, originally introduced in the context of Noise Contrastive Estimation and later popularized in representation learning frameworks such as CPC and SimCLR[1,2]. Given a batch of sequence embeddings $\boldsymbol{z}_i^{seq}$ and signal embeddings $\boldsymbol{z}_j^{sig}$, we compute a similarity matrix:

$$S_{ij} = \frac{\boldsymbol{z}_i^{seq} \cdot \boldsymbol{z}_j^{sig}}{\tau}$$

where $\tau$ is a temperature parameter controlling distribution sharpness. For each sample $i$, the matched sequence-signal pair $(i, i)$ is treated as the positive pair, while all other batch elements serve as negatives. The symmetric InfoNCE objective optimizes:

$$\mathcal{L}_{InfoNCE} = \frac{1}{2}\left(CE(S, diag) + CE(S^T, diag)\right)$$

where $CE$ denotes cross-entropy and $diag$ indicates the matching index. This formulation is similar to the alignment strategy used in CLIP-style multimodal training[3]. InfoNCE enforces instance-level alignment but does not explicitly account for label structure.

*Label-aware contrastive loss.* Because DNA damage detection is a supervised task with explicit lesion labels, we further implemented a label-aware contrastive loss, in which positives are defined as all samples sharing the same damage label, and negatives are those with different labels. This formulation is inspired by supervised contrastive learning[4], which extends InfoNCE by grouping samples based on class identity rather than strict instance matching. Similarly, we first compute the similarity matrix $S_{ij}$ as before. Let $y_i \in \{0,1\}$ denote the binary damage label. For each anchor $i$, we define the set of positives $\mathcal{P}(i) = \{j \neq i \mid y_j = y_i\}$ and the set of all candidates excluding self-pair $\mathcal{A}(i) = \{j \neq i\}$. The label-aware contrastive loss for anchor $i$ is:

$$\mathcal{L}_i^{label} = -log\, \frac{\sum_{j \in \mathcal{P}(i)} exp\,(S_{ij})}{\sum_{k \in \mathcal{A}(i)} exp\,(S_{ik})}$$

The total loss is averaged over anchors with at least one positive:

$$\mathcal{L}_{label} = \frac{1}{|\mathcal{V}|} \sum_{i \in \mathcal{V}} \mathcal{L}_i^{label}$$

where $\mathcal{V} = \{i \,|\, |\mathcal{P}(i)| > 0\}$. Under this objective, each embedding is encouraged to align with embeddings from the same damage class across modalities while being separated from embeddings of different classes.

*Damage-Focused Contrastive Loss.* Given the class imbalance in genome-wide lesion detection, we restrict contrastive alignment to damaged samples only. Let $D = \{i | y_i = 1\}$ be the index set of damaged samples in a minibatch and let $M = |D|$. If $M < 2$, we set $\mathcal{L}_{\text{damage}} = 0$. Otherwise, we form the damaged-only similarity matrix $S^{(D)} \in \mathbb{R}^{m \times m}$ by selecting the sequence and signal embeddings of samples in $D$ and computing $S_{ab}^{(D)}$ as before. We then apply a symmetric InfoNCE objective restricted to the damaged subset[5]:

$$\mathcal{L}_{damage} = \frac{1}{2}(CE(S^{(D)}, diag) + CE((S^{(D)})^{\top}, diag))$$

where $diag$ indicates the matched sequence-signal pairs within the damaged subset. Equivalently, for an anchor $a \in \{1, \dots, M\}$, we have:

$$\mathcal{L}_a^{damage} = -\,log \frac{exp\left(S_{aa}^{(D)}\right)}{\sum_{b=1}^{M} exp\,(S_{ab}^{(D)})}$$

The total loss is averaged over anchors:

$$\mathcal{L}_{\text{damage}} = \frac{1}{M}\sum_{a=1}^{M} \mathcal{L}_a^{\text{damage}}$$

This objective strengthens alignment between matched sequence-signal representations only for lesion-containing instances, encouraging a compact and discriminative embedding structure for the biologically informative minority class while avoiding unnecessary constraints from abundant undamaged samples.
